# Temporal variation in demography of temperate bats: consequences for population dynamics and disease

**DOI:** 10.64898/2026.02.25.707526

**Authors:** Susannah Gold, Simon Croft, Richard Budgey, James Aegerter

## Abstract

Bat populations experience inter-annual variation in demographic rates in response to environmental conditions. This variation has the potential to impact population sizes and structures, in addition to population-level processes such as disease spread. To establish the influence of variation in demography on these processes, we develop a spatial, individual based model of a serotine bat (*Eptesicus serotinus)* population, within which we introduce a synthetic lyssavirus-like disease. Model results show that increasing demographic variation, particularly in survival rates, may drive substantial population decline in bat populations. Increasing environmental fluctuations driven by climate change may therefore be problematic for population persistence. The likelihood of disease persistence was also reduced by increasing variation. These findings highlight the limitations of only considering mean demographic rates for prediction of population size change and disease dynamics from models.

## Introduction

Estimating demographic parameters, such as survival and reproductive rates, is central to predicting future population trends for wildlife. These parameters also inform population models underlying epidemiological models for disease in wildlife. Within these models, demographic processes are often simulated based on mean annual rates, which are assumed to be fixed across time. However, for many species, demography is highly influenced by the external environment, leading to temporal variation in survival and reproductive rates. This stochasticity can have important impacts for population growth rates and persistence (Boyce et al. 2006). In temperate bat species, variation in populations between years, both in terms of observed abundance and demographic rates, has been regularly reported (Ransome and McOwat 1994; Frick et al. 2010; Fleischer et al. 2017; Reusch et al. 2019). Understanding the impact of this variation on the demography of temperate bats is of substantial interest both from a conservation and public health perspective. Bat species are host to a number of pathogens with zoonotic potential (Tan et al. 2023), and spill-over risk may be influenced by demographic change in the reservoir hosts (Plowright et al. 2017). Predicting future trends in populations of bats, and their pathogens, therefore requires an understanding both of the influence of current temporal variation in demographic parameters, and of future variation, which may increase with climate change.

The impact of environmental variation on population and disease outcomes will depend on the specific species’ lifestyle (Reusch et al. 2019). Bats in general demonstrate a slow (K-selected) life-history, with long lifespans and low reproductive rates, particularly relative to other small-bodied mammals. As a result, populations are expected to be particularly sensitive to reductions in adult survival. High mortality within years may result from extreme weather events, such as heatwaves or harsh winters, which may increase with climate change (Salinas-Ramos et al. 2023; Reusch et al. 2019). Reproduction in temperate bats is also vulnerable to environmental conditions, particularly as there is a relatively short window during spring and summer in which gestation, lactation, and growth of juveniles to independence occurs. As bats are income breeders, which rely on concurrent foraging to provide energy for reproduction, resource availability during this window is central to successful reproduction. As a result, environmental factors that reduce insect populations or increase energy requirements (e.g., cold weather) during this period can substantially impact on reproductive success, for example leading to failure of pregnancies or abandonment of pups due to insufficient resource. Understanding the specific environmental drivers of demographic change in bat populations is challenging, requiring long-term capture-mark-recapture projects (Frederiksen et al. 2014). Here, we do not directly consider the causes of inter-annual variation but instead explore the potential outcomes of this variation for a bat-disease system.

Within bat populations, the influence of environmental variation is expected to vary between different demographic classes. Males, and non-breeding females, avoid the energetic costs of pregnancy and lactation faced by breeding females, therefore potentially reducing the impact of resource limitation during spring. Juveniles in general are expected to have lower baseline survival than adults (Reusch et al. 2019) and may be particularly vulnerable to poor environmental conditions soon after independence. An outcome of these demographic differences between classes is that population structure may vary over time, with potential implications for disease processes. For example, younger bats are more likely to be susceptible to disease due to lack of prior exposure and development of immunity, whereas breeding females may have higher contact rates due to their use of large maternity roosts. Demographic shifts in populations may therefore have implications for disease processes that are not attributable solely to population densities.

We develop a simulation model to test the potential impact of inter-annual variation in demographic parameters, using the case study of the serotine bat (*Eptesicus serotinus)*. This species is of interest for multiple reasons. It is currently listed as vulnerable by the IUCN in Great Britain and is limited in distribution to Southern England and Wales. While monitoring suggests that the population is currently stable (BCT 2023), climate change could be expected to benefit this species, which is widespread in warmer European climates. However, the increasing variability in weather patterns that is predicted to occur could have negative impacts. Serotine populations are also of interest as they are considered the primary reservoir for European Bat Lyssavirus-1 (EBLV-1), which was first detected in the UK in 2018 (Folly et al. 2021). An improved understanding of the population dynamics of this species, and how it may react in response to future environmental change, will inform future efforts to understand the dynamics of viral pathogens within this species.

The model developed is a spatially explicit, individual-based model parameterised to realistically represent serotine bat demography and spatial structure. Demographic rate parameters are based on a long-term capture-mark-recapture study at a serotine roost, which provides detailed information on how these rates vary between classes and across years. Within this framework, we introduce a synthetic lyssavirus-like disease to test interactions between demography and epidemiology. We use this model to explore two key questions: i) how does increasing inter-annual variation in demographic parameters influence predictions of future population size change? and ii) does inter-annual variation influence the likelihood of disease persistence within this population?

## Methods

In this paper, we describe the development and calibration of an individual-based model of a serotine bat population. This model is parameterised to produce plausible dynamics which reflect the current UK serotine population. The parametrisation of the simulated bat population is primarily drawn from a single long-term study of serotine bats at one site, and as such, represents a coherent and robust ecological model of how we think the population dynamics of this species worked through that period. Using this model, we then test the influence of increasing the level of variation that occurs between years in demographic parameters on the predicted change in population size, and the predicted disease dynamics.

### Model summary

A full model description following the ODD protocol (Overview, Design and Details) (Grimm et al. 2020) is provided in the supplementary material (Appendix S1). In brief, the model simulates the complete life-histories of individual bats as they develop through a series of demographic classes (pup, juvenile, immature, reproductively mature). The model runs through an annual cycle in monthly time-steps, allowing for season-specific behaviours and demographic rates. We assume births occur simultaneously in June and pups become independent juveniles in July. Bats are assumed to hibernate between October-March, with an active period from April-September.

Bats are divided into social communities of variable size within a model arena of 30km by 30km (∼5,000 bats total across 45 communities). Our model landscape is a mosaic of cells constructed using an irregular geometry, with each patch hosting a community of serotine bats. Communities are assumed to occupy an exclusive geographic footprint consisting of a single maternity roost, where pups are produced by reproductive females, and a network of “satellite roosts”, used by all demographic classes at varying times, but which males and a proportion of non-breeding females use through the active period. The location of bats (maternity-roost or satellite-roost) is used to define contact rates for disease transmission. For simplicity, in this study we have assumed that there is no spatial variation in demographic parameters, with all communities having the same survival and reproductive rates.

The potential for an extended pre-reproductive period in females, as previously observed for serotine bats (Chauevenet et al. 2014) was modelled by stochastically determining whether females became reproductively mature, with differing probabilities for juvenile and adult females. Once an individual was reproductively mature, it was assumed it would attempt to reproduce in every following year, with success stochastically determined based on a fecundity parameter. Serotine bats in general produce only a single offspring, but it was assumed there was a low probability of twinning.

Limited information is available on dispersal in bats. Genetic evidence suggests dispersal is male-biased, with the majority of females showing natal philopatry (Moussy et al. 2015). It was therefore assumed all juvenile males would disperse from their maternity roost in their first year, in addition to a small, fixed proportion of non-breeding females (2%). Given that recorded dispersal distances are high relative to the arena size (Bogdanowicz et al. 2013; Martinoli et al. 2020), we assumed a new community within the arena was selected randomly.

In each month, survival of bats was determined with probabilities based on four factors: age, breeding status, season, and year “quality”. Pup survival (birth to weaning) was dependent on maternal survival and an additional probability of pup death. Following weaning, juveniles were assumed to have higher mortality than adult bats. For females, breeding was assumed to result in a survival penalty in the breeding period (June-July) due to the high energetic costs of reproduction. In the absence of evidence to describe the mortality of adult males, we assume their survivorship follows that of non-breeding adult females, based on the assumption that they similarly do not face the energetic costs of pregnancy and lactation. Mortality rates in all demographic classes were assumed to be lowest during the hibernation period, due to the lower risks of predation and reduced energetic requirements (O’Shea et al. 2011; Reusch et al. 2019).

### Model fitting

Parameterisation of the base model was calibrated to produce a stable population with realistic demographic behaviour in the absence of disease. For each of the demographic parameters (Table 1), plausible ranges were extracted from a capture-mark-recapture (CMR) analysis of a serotine population at a single site (see Appendix S1), and from the literature (Harbusch and Racey 2006). The CMR dataset has previously been analysed in Chauvenet et al. (2014). We conducted a re-analysis of this data to estimate survival parameters (divided into juveniles, non-breeding adults, and breeding females) and reproductive parameters (primiparity for juveniles, primiparity for previous non-breeders and fecundity), both as mean annual estimates, and to estimate inter-annual variability. The 95% interval for mean annual estimates was used as the plausible range for these parameters. For parameters that it was not possible to estimate using this dataset, either literature estimates were used to generate plausible ranges (e.g., pup survival), or wide plausible ranges were assumed (e.g., proportion of non-breeding females within maternity roosts). Within these ranges, 1,500 different parameter combinations were generated using Latin hypercube-sampling. For each parameter combination the model was run for 30 years, and the parameter combination was chosen that produced dynamics closest to empirically observed patterns. Plausible dynamics were defined as those that produced patterns consistent with observations from UK and European Serotine populations. These patterns were: i) stable population size, ii) population sex ratio and iii) the proportion of females in maternity roosts that are reproductively active.

**Table 1:** Calibrated parameter values for serotine bat demography.

| Parameter | Value | Uncertainty range | Units |
| --- | --- | --- | --- |
| Juvenile primiparity | 0.12 | 0.05-0.26 | Probability per year of juvenile female of becoming reproductively mature |
| Non-breeder primiparity | 0.24 | 0.20-0.39 | Probability per year of non-breeding adult of becoming reproductively mature |
| Fecundity | 0.93 | 0.84-0.97 | Probability per year of reproductively mature adult producing a pup |
| Twinning rate | 0.02 | 0-0.05 | Probability of litter size = 2 |
| Pup survival | 0.94 | 0.7-1 | Probability per pup of surviving from birth to independence |
| Juvenile survival | 0.53 | 0.48-0.83 | Survivorship between independence and end of first hibernation |
| Adult survival | 0.91 | 0.69-0.93 | Annual survival probability for males, and female non-breeders (inc. immatures) |
| Breeder survival | 0.83 | 0.70-0.90 | Annual survival probability for reproductively mature females |
| Proportion non-breeders at maternity roost | 0.68 | 0.05-0.95 | Annual probability that non-breeder will be present in maternity roost, rather than satellite roost |

Population change was matched against information from the Bat Conservation Trust (BCT) National Bat Monitoring Programme (NBMP). Roost counts suggest that UK Serotine bat populations are approximately stable, with the 95% confidence interval for annual change containing 0 (-1.5% to 1.8%). Therefore, for each simulation, the mean population size change between years was calculated as a summary statistic and compared against a change of 0.

Population-level sex ratios were extracted from studies reporting on serotine bat submissions to disease-testing programmes. The majority of studies on serotines have been focused on maternity roosts, where adult males are excluded. By contrast, passive surveillance schemes should be relatively unbiased samples of the total population. Across 6 sources (from 5 countries) identified which reported sex of submitted bats (1,559 bats total), an overall male-biased sex ratio of 60-40 was found (Van der Poel et al. 2005; Muhldorfer et al. 2011; Harris et al. 2006; Picard-Meyer et al. 2016; APHA Bat Rabies Dashboard (accessed 8/3/2023); Orlowska et al. 2020). Proportion of males at the end of each 30-year simulation was extracted and compared against the empirical estimate of 0.6.

For the proportion of breeding females at maternity roosts, in a study at maternity roosts, Catto (1996) reported that of adult females captured, 66% were reproductively active when excluding those of undetermined status. This estimate aligns with the CMR dataset from the UK, where 67% of unique annual captures of adult females were assigned as reproductively active when excluding unknown status. The mean proportion of females marked as present in maternity roosts during the breeding season that were reproductively active was therefore extracted and compared to a value of 0.66.

For each simulation, Euclidean distances were calculated between the summary statistics and empirical estimates, and the parameter combination producing the minimum distance selected. These fitted parameter values are presented in Table 1. The model was then run for 100 simulations for this parameter combination and the distribution of outputs plotted to visualise model fit (Appendix S2: Figure S1).

### Disease scenarios

Following establishment of a stable population model, we introduced a simulated disease into the population to explore how epidemiology may be influenced by inter-annual demographic variation. In this study our pathogen deliberately resembles *Lyssavirus hamburg* (hereafter EBLV-1), although because there remains considerable uncertainty around the pathology of lyssaviruses in bats (see Gentles et al. 2020 for a review of bat-lyssavirus models) we do not attempt to explicitly replicate EBLV-1 in this system, but rather produce a simplified pathology to demonstrate the interactions between demography and epidemiology.

Pathology was simulated as a SEIR process (Susceptible, Exposed/Latent, Infectious, Recovered/Immune). Following exposure, bats entered the latent phase, the duration of which was randomly sampled from a Poisson distribution. Based on evidence from pathology of rabies in bats (Davis et al. 2016) it was assumed pathology was “paused” during hibernation, therefore progression through the latent period only occurred during active months. At the end of the latent period, bats either became infectious and died, or experienced a sub-clinical infection and became immune, with the duration of immunity also sampled from a Poisson distribution, after which bats returned to being susceptible.

Contact rate between bats were assumed to depend on roost type, with three scenarios; contact within a maternity roost, contact within a satellite roost within the same community, or with much lower probability, contact between communities. In common with many temperate bat species, potentially large numbers of bats aggregate within maternity roosts, whereas in satellite roosts, bats are expected to be roosting singly or in small groups, with infrequent contact. Some demographic classes are assumed to be excluded from maternity roosts (e.g., adult males), but others might move between maternity and satellite roosts (e.g., non-reproductive females). A transmission function was used that was intermediate between density and frequency-dependent, with contact rates increasing with the number of individuals within the community but plateauing as this population size increased (Figure 1). Transmission of disease between roosts could occur either through dispersal movements of latently infected juveniles, or throughout the year through inter-community contacts between males and non-breeders in satellite roosts. Full details are provided in the ODD (Appendix S1). Parameters used are shown in Table 2.

**Figure 1:**
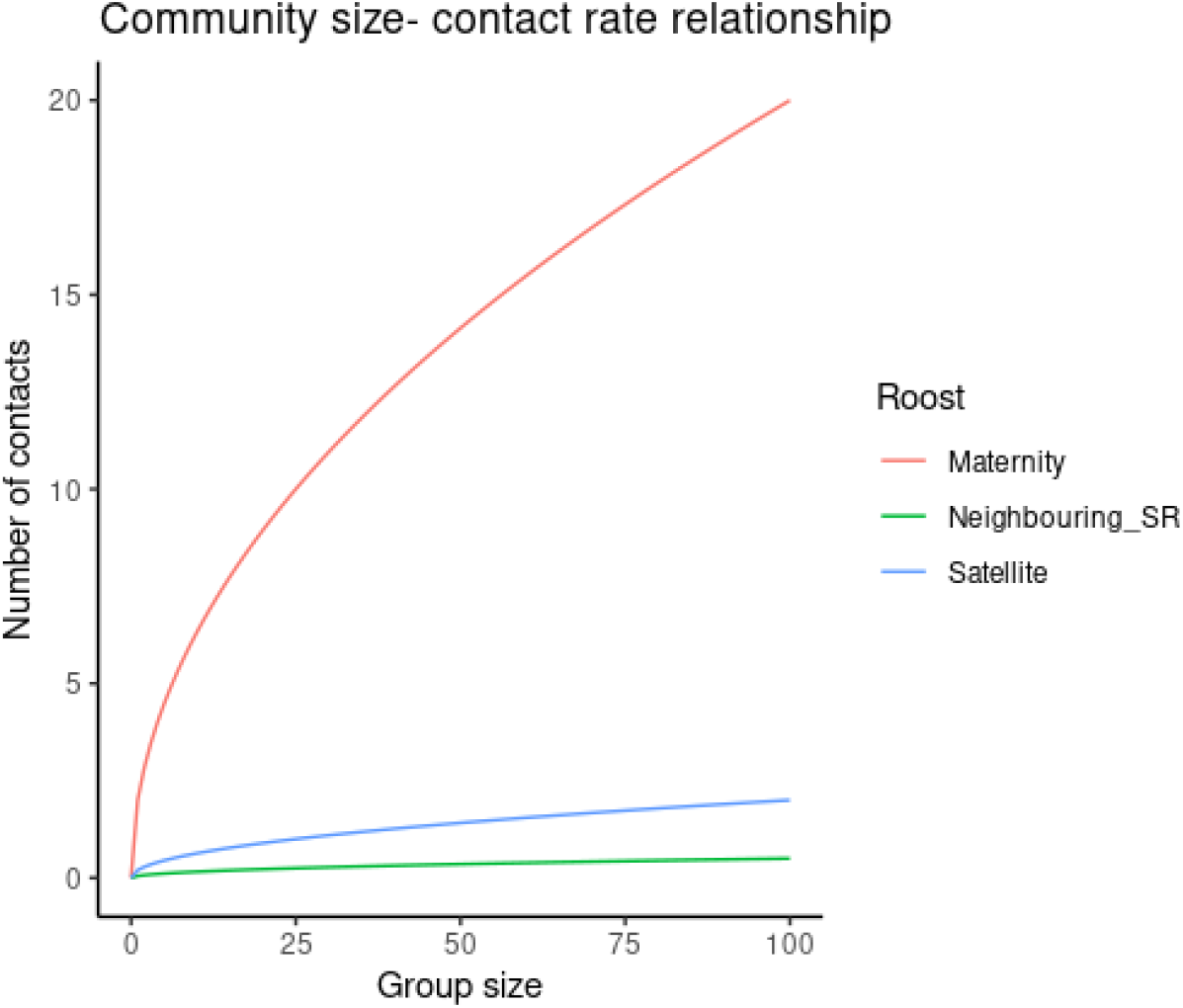
Assumed relationship between group sizes (for maternity roosts, satellite roosts, and neighbouring satellite roosts respectively) and contact rate between individuals.

**Table 2:** Assumed disease parameters for synthetic disease.

| Parameter | Value | Unit | Justification |
| --- | --- | --- | --- |
| **Transmission probability** | 0.75 | Probability of disease transmission per contact | Assumption that disease highly transmissible |
| **Probability of fatal infection** | 0.15 | Probability that infection leads to mortality | Low probability of clinical infection based on observations from bat-lyssavirus systems (George et al. 2011; Blackwood et al. 2013; Kim et al. 2023) |
| **Duration latency** | 1 | Mean duration of latent period, drawn from Poisson distribution | Assumed mean latent period ~ 1 month, with potential for longer durations, in line with other bat-lyssavirus models (George et al. 2011; Blackwood et al. 2013; Kim et al. 2023) |
| **Duration infectious period** | 1 | Maximum duration of infectious period (months) | Assumed duration of infection was short relative to time step of 1 month |
| **Duration immunity** | 6 | Mean duration of immunity period drawn from Poisson distribution (months) | Assumed variable duration of temporary immunity |
| **Maternity roost contact rate** | 2.5 | Contact rate per month in maternity roost | Assumption of high contact rates within dense maternity roosts |
| **Satellite roost contact rate** | 0.25 | Contact rate per month in satellite roost | Assumption of low contact rates in satellite roosts, as usually found singly or in small groups |
| **Neighbouring satellite roost contact rate** | 0.05 | Contact rate per month in satellite roost with neighbouring communities | Assumption of very low contact rates between neighbouring communities |

### Annual variation model scenarios

The base model used mean annual estimates to determine reproduction and survival, which were calibrated to produce plausible population dynamics, with a low-level of disease. From this baseline, we tested different scenarios for increasing levels of inter-annual variation. Initially, we assumed all parameters co-varied, so that a “good” year increased both survival and likelihood of reproduction (primiparity and fecundity). It was assumed that all variation is environmentally driven, as opposed to other external factors, such as predation. This variation was introduced by, for each model year (January-December), selecting a “quality” from a normal distribution (Q∼N(0,σ)), with positive values reflecting a “good” year and negative values reflecting a “bad” year. The greater the variation in this distribution (σ), the higher the level of inter-annual variation. Survival and reproductive parameters were input as logit values. This quality for a particular year was used to modify the reproductive and survival parameters (Py) using the following relationship:

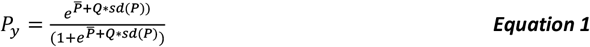

Where the mean annual rate 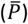 and level of inter-annual variation (*sd* (*P*)) are represented on the logit scale. For juvenile survival, non-breeder survival, and breeder survival, it was possible to extract an estimate of the inter-annual variability (σ) from the CMR analysis. No information on inter-annual variability on other parameters was available, therefore arbitrary values were assumed (σ=1). Illustrative examples of the resulting probability distributions are shown in Figure 2 across increasing levels of inter-annual variation.

**Figure 2:**
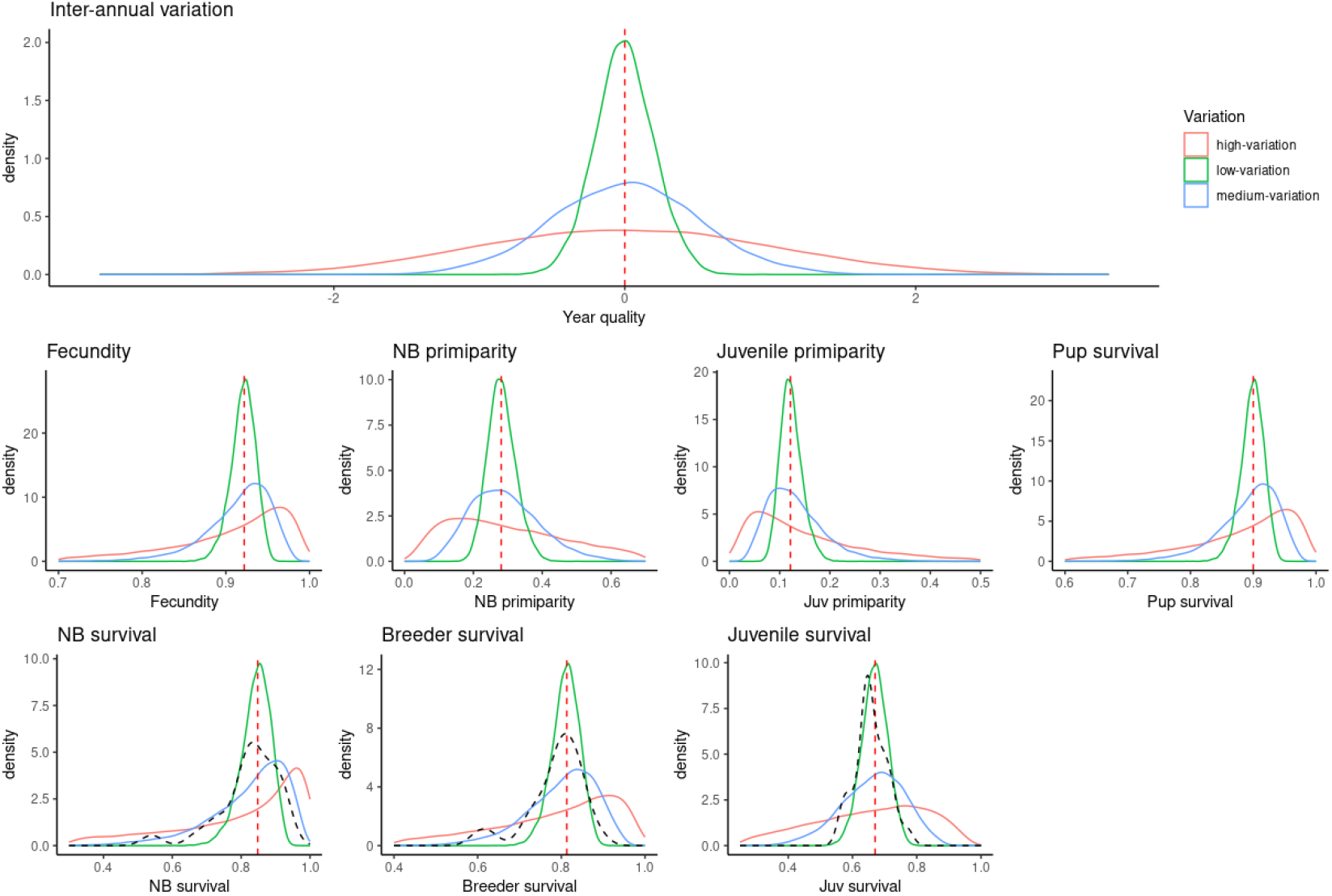
Example parameter distributions for different degrees of inter-annual variation in demographic parameters. The red dotted line indicates the mean fitted value (no inter-annual variation). Distributions shown by black dotted lines reflect estimated distributions from CMR analysis. Red vertical line indicates σ of 0, green=0.2, blue=0.5 and red=1.

We tested the influence of increasing inter-annual variation (with σ increasing from 0 to 1, in intervals of 0.2). In each case, the median parameter values remain constant, but variation around this increases (Figure 2). For each scenario, following a 30-year burn-in to allow population dynamics to stabilise, we ran 50 simulations without disease, and extracted the predicted change in population size across a 20-year simulation. To test the epidemiological implications, we then introduced disease and extracted the time from disease introduction to disease extinction (persistence duration), as well as the population-level seroprevalence and number of infected bats.

To test the key drivers of demographic and disease change, after co-varying all parameters, we separated parameters into those defining survival of free-flying individuals (juvenile survival, breeder survival, and non-breeder survival) and those influencing reproduction (primiparity, fecundity, and pup survival). Simulations where all parameters were allowed to vary independently are also presented in the supplementary material (Appendix S1).

### Demographic and community level impact

To explore in more detail how variation in year qualities influences population and disease processes, in addition to running simulations where the quality of each year was randomly selected, we also ran a specific pattern of years, alternating between “good” (Q=1), “bad” (Q=-1) and average (Q=0) years for 10 years for 10 simulations. To illustrate the impact on demography, we visually represent how the population size, proportion of individuals reproducing and number of newly produced juveniles varies between years. From a disease perspective, we show how prevalence and seroprevalence vary, both at a population and community-level.

### Sensitivity analysis

A global sensitivity analysis was run to test which parameters are most influential for demography and disease dynamics. A total of 250 parameter combinations for all demographic parameters were selected using Latin hypercube sampling (see Appendix S1 for ranges). The model was run for 20 years with the synthetic disease included and population change, and duration of disease persistence extracted. Partial rank correlation coefficients were calculated to visualise the relationship between input parameters and output parameters.

## Results

### Predicted population change under increasing inter-annual variation

Increasing inter-annual variation in demographic parameters substantially influenced the predicted change in population size over time. The model was calibrated so that using fitted parameter values with no inter-annual variation, the population size remained approximately stable. As randomly sampled inter-annual variation increased from this baseline the predicted outcomes for populations became more variable between simulations. While in some simulations, high variation led to substantial population growth, it also increased the probability of population crashes, with at the highest level of variation considered (σ=1), 30% of simulated populations declined to less than 5% of the starting population. The general trend was for a substantial decrease in serotine population size over time as the variability increased (Figure 3), this trend was consistent whether all parameters co-varied (Figure 3) or varied independently (Appendix S2: Figure S2).

**Figure 3:**
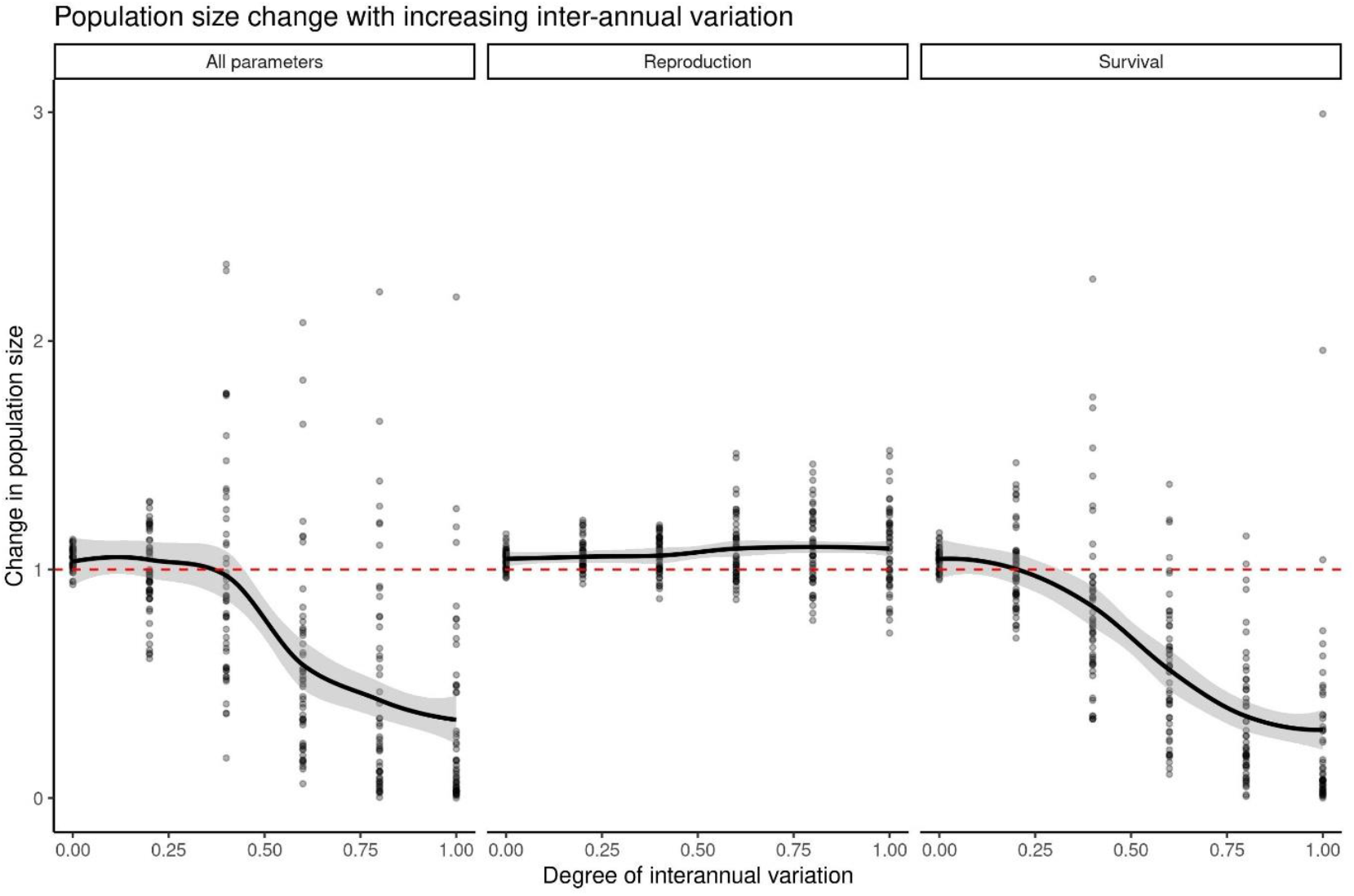
Relationship between degree of inter-annual variation and population size change over 20 years. Population size change is defined as the total population at the end of the simulation, divided by the start population for that particular simulation.

When variation was limited to either reproductive or survival parameters, inter-annual variation in survival was a more influential driver of population change. Where only reproduction was varied, although variation in population size increased between simulations, there was little impact on mean population size change. This result suggests that serotine bat populations may be able to buffer variation in reproductive success, as long as survival remains high.

### Persistence of disease under increasing inter-annual variation

The level of inter-annual variation also influenced disease dynamics. Our synthetic disease was seeded into 20 bats in the stable burn-in population. In the no variation scenario, on average disease persisted for 9 years post-introduction (Figure 4), with 18% of simulations having disease persistence to the end of the simulation (20 years). Increasing variation in demographic parameters decreased the mean duration for which disease persisted, to 5.5 years when all parameters were allowed to co-vary, with only 4.1% of simulations having disease persistence across 20 years. As with population change, this reduction was driven primarily by the variation in survival parameters, with only a limited reduction in duration of persistence when only reproduction varied. While disease persistence was negatively impacted by increasing variation in demogaphic parameters, the total number of infectious bats across the course of the simulation remained relatively stable (Appendix S2: Figure S2). While disease persisted for a shorter period, in some simulations variability led to high numbers of infected bats, resulting in similar averages.

**Figure 4:**
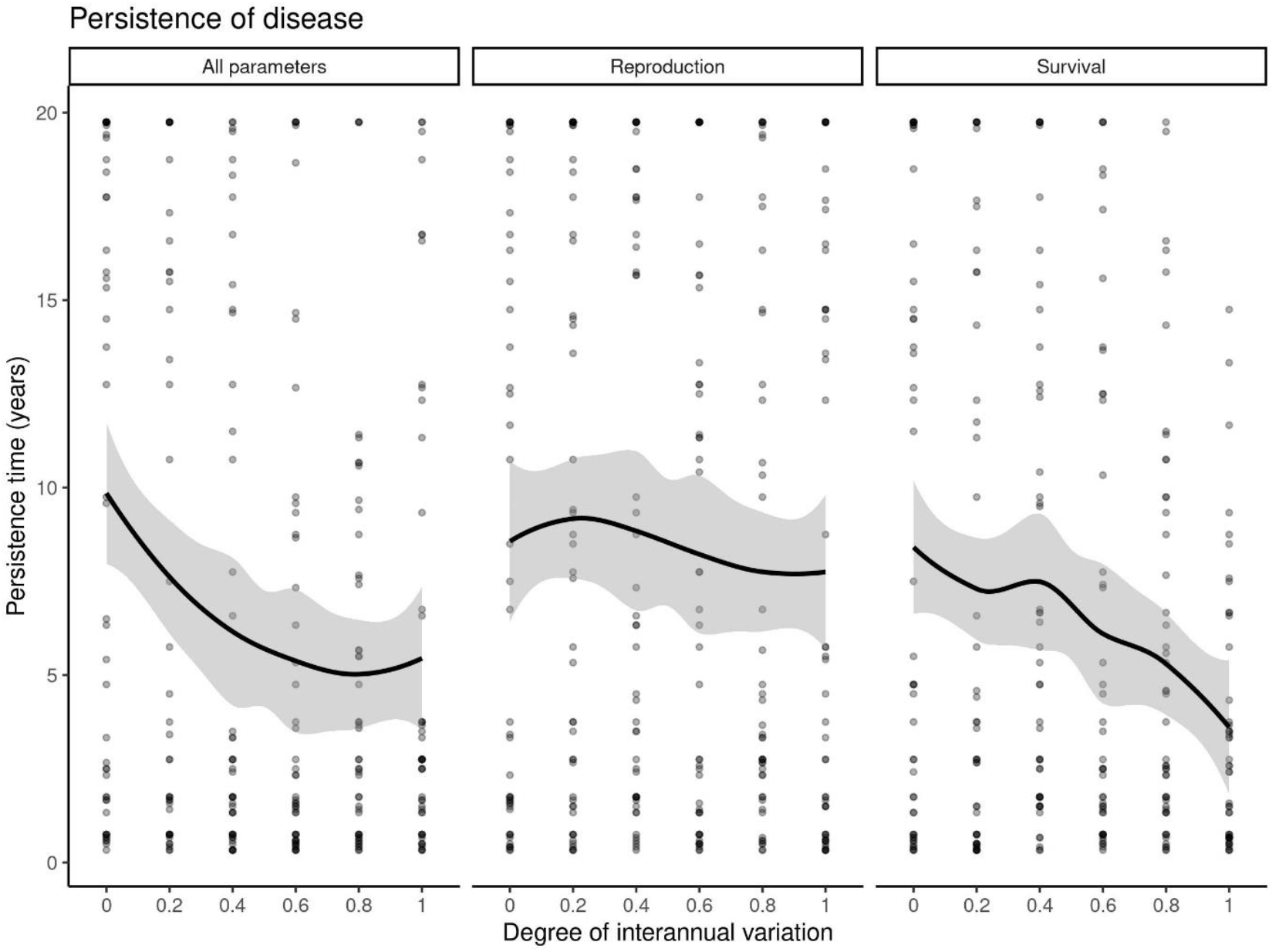
Relationship between degree of inter-annual variation and time to disease fade-out (Persistence). Three scenarios were considered: all parameters co-vary, variation in reproductive parameters only (primiparity, fecundity and pup survival) and only survival parameters co-vary (juvenile, non-breeder, and breeder survival). Trend lines were generated using the loess function from the geom_smooth function in R.

### Changes in demographic structure with change in quality

To explore the relationships between year quality, demography and disease dynamics, the model was run for a pre-specified pattern of “good”, “bad” and “average” years (Figure 5). Within years, populations fluctuate in size, due to the birth pulse in summer, and differing mortality rates between hibernation and the active period. In good years, the improved reproductive success increased the birth pulse, with more individuals present in maternity roosts and a higher number of juveniles produced. Lower mortality led to a slower reduction in population size.

**Figure 5:**
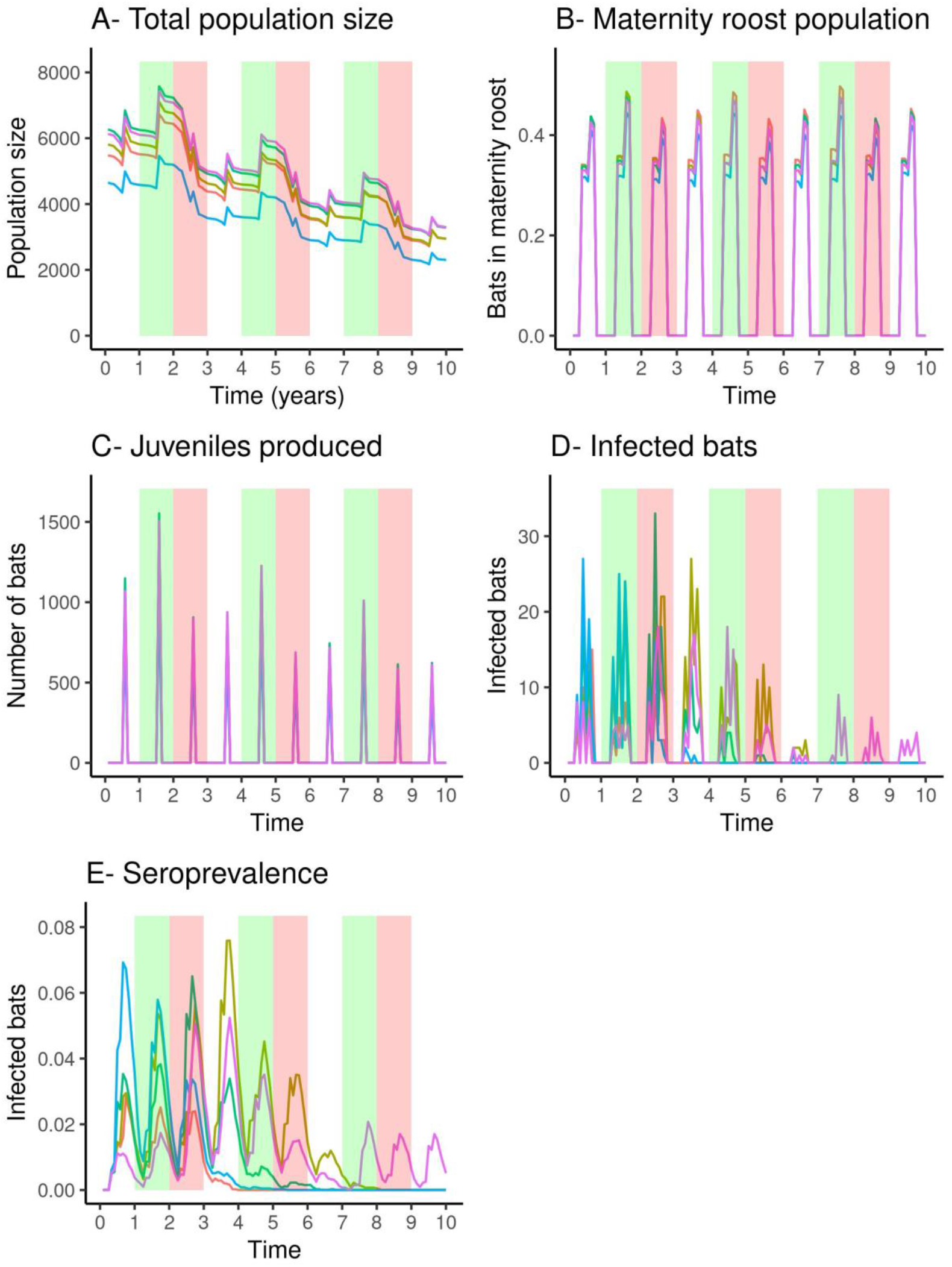
Outputs from individual simulations showing impact of “good” years (Green-Quality=1), “bad” years (Red-Quality=-1), and “average” years (White-Quality=0) on population structure (A-C) and disease dynamics (D-E). Each colour represents an individual simulation.

While the higher density of individuals in maternity roosts provides more contact opportunities, and there is a greater influx of new susceptible individuals in good years, within a single simulation this did not have obvious impacts on the disease dynamics. Disease processes showed high levels of variation between simulations. As shown in Figure 6 for a single simulation, disease did not persist in any one population throughout the simulation but was maintained at a community-level. Seroprevalence (the immune proportion of each community) varied substantially between communities across the course of the simulation. The persistence and number of cases is driven by stochastic processes, such as the likelihood of dispersal of a latently infected individual to a new community, or inter-community contact.

**Figure 6:**
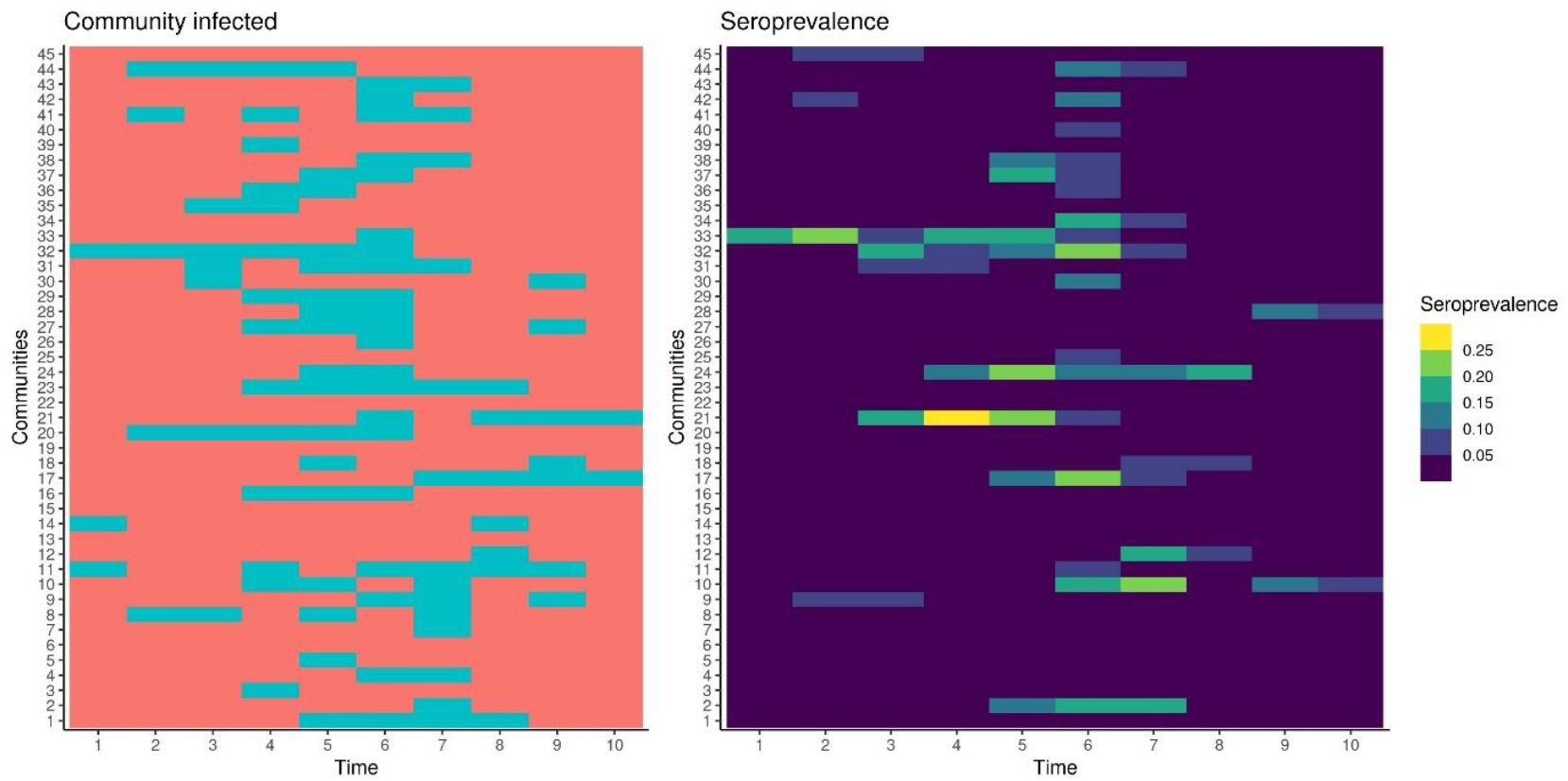
Persistence of disease at a metapopulation level for a single simulation. A) shows which of the 45 total communities contain at least one infected individual in each year of the simulation (blue= infected, red=uninfected). B) shows the average seroprevalence across each year of the simulation for each community.

### Sensitivity analysis

A sensitivity analysis was conducted to explore how influential uncertainty in demographic parameters was for population change and persistence of the synthetic disease within the model (Figure 7). For population size change, model outputs were highly sensitive to uncertainty in survival parameters, in particular adult survival, as expected for a long-lived species like the serotine bat. The uncertainty in reproductive parameters (primiparity and fecundity), was less influential. Twinning and pup survival had very little influence.

**Figure 7:**
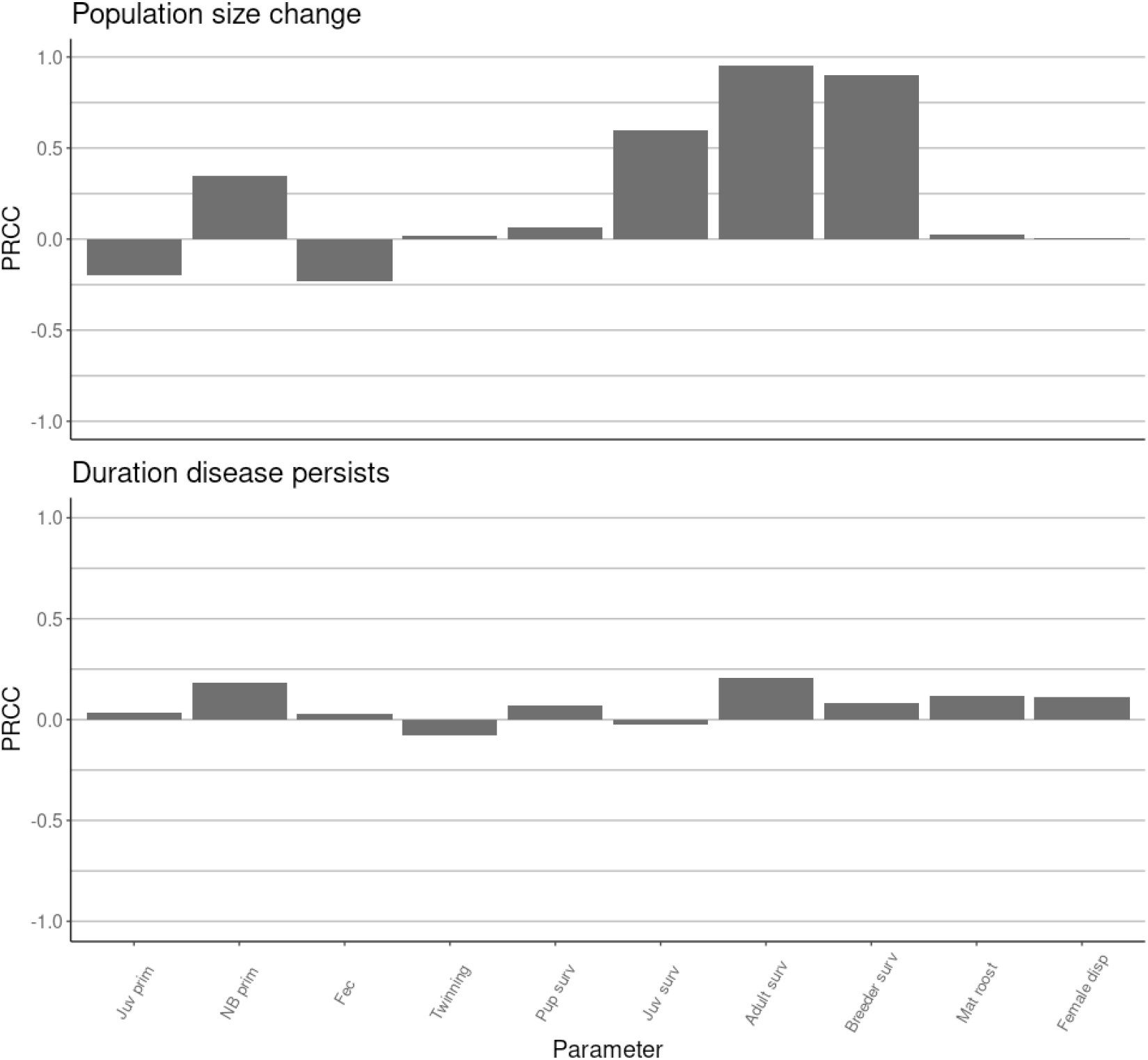
Global sensitivity analysis using LHS-PRCC method. Bars indicate relationship between input parameters (Juvenile primiparity, Non-breeder primiparity, Fecundity, Twinning, Pup survival, Juvenile survival, Adult survival, Breeder survival, Proportion of non-breeding females using maternity roost and Female dispersal) and A) the size in overall population over 20 years and B) the duration for which disease persisted.

For disease persistence, no demographic parameters were highly influential, supporting earlier results that there are high levels of stochasticity which determine the differences between simulations. Adult survival was the most influential parameter for persistence as well as population size. Following this, uncertainty in non-breeder primiparity was also influential, with a higher rate of transition to breeders increasing disease persistence. It was assumed that once females transition to being breeders, they move into the maternity roost, with higher inter-individual contact rates. Therefore, higher transition probabilities may increase the force of infection.

## Discussion

Environmental variation can be highly influential for population processes. Using a detailed individual-based model, we found that variation in demographic rates in a temperate bat population could have substantial impacts for predictions of future population sizes, and potentially for disease dynamics. As expected for a long-lived species like the serotine bat, variation in adult survival in particular was the primary driver of population processes. As variation in survival increased, in general population sizes tended to decrease. This highlights the potential threat posed to bat populations by significant mortality events, which may be caused by extreme weather. While “good” years may have survival approaching 100% and high levels of reproduction, this does not counteract the impact of a “bad” year where a substantial proportion of the breeding population is lost, due to the persisting impact of the loss of breeding individuals (Fleischer et al. 2017).

In contrast to survival, variation in reproductive rates had little impact on population dynamics. We assumed that in “good” years, a higher proportion of juvenile and adult non-breeding bats would transition to become breeders, and that breeders would have greater success in raising pups to the point of independence. However, even in good years, population growth rates in serotine bats are constrained by the ability to only have one (or in rare occasions, two) pups in a year. This low reproductive rate is buffered by the longevity of the species as individuals can afford to have a delayed onset of reproduction and produce only one pup per year if they have a long-life as a breeding individual. Increasing reproductive rates leads to a trade-off due the high investment required, with breeding females having higher mortality rates than non-breeding adults. The population increases driven by higher proportions of individuals choosing to breed, are therefore partially counteracted by the resultant higher mortality in breeding individuals.

In addition to population growth impacts, demographic variation also influenced the persistence of the synthetic “lyssavirus-like” disease. Increasing variation leads to greater potential for stochastic fade-out of disease in poor years, where there are smaller populations, leading to lower contact rates. However, the predicted mean total number of infectious bats was relatively robust to increasing variation, with predictions biased upwards by some simulations with much higher numbers of bats becoming infected. These results highlight the importance of considering inter-annual variation for predicting disease dynamics. Dynamics are substantially different, despite demographic parameters having the same median values. In addition, the outcome of disease introduction was highly variable between simulations. This variation arises partially due to the spatial structure in serotine populations. For disease to persist, it needs to be able to move between communities to “escape” communities with high seroprevalences. This fits with empirical observations of bat-lyssavirus dynamics, where substantial fluctuations in community-level seroprevalence have been observed (e.g., for *Myotis myotis*, Pons-Salort et al., 2014). While little information is known about serotine movements, it is expected that outside of maternity roosts, contact rates between individuals are relatively low. These inter-community transmission events are therefore dependent on presumably rare events such as juvenile dispersal of latently infected individuals, or inter-community contact with an infected individual.

Other factors driving disease persistence include the assumption of extended latent periods allowing disease to persist through periods of low contact rates in the winter. Disease can then be amplified by high contact rates in maternity roosts in the summer. This is observed in the model with peak numbers of infected bats occurring in the summer period. Break-up of maternity roosts into smaller satellite roosts, and mixing of males and females in the autumn, then potentially allows for further spatial spread prior to hibernation. While we did not aim to have specifically modelled realistic EBLV-1 dynamics, this provides evidence that incursions of pathogens, including lyssaviruses, into UK bat species may be able to persist long-term, despite the fluctuations in populations that occur in this range-edge population. However, the impact of population demography on disease is highly dependent on the assumed transmission mechanism. While we aimed to produce plausible dynamics, contact rates and transmission functions were assumed. If contact rates are more linearly dependent on density, this could increase the sensitivity to population fluctuations (Borremans et al. 2017), and vice-versa if transmission is frequency dependent. Collecting contact rate data on bats is highly challenging. While out of the scope of this paper, future work could aim to improve the realism of the synthetic disease by fitting the disease model to observed patterns of prevalence and seroprevalence from serotine populations.

There are a number of future potential uses of the model developed. Ideally, demographic parameter estimates could be linked directly to weather patterns, allowing predictions to be tied to future climate forecasts (e.g., Lee et al. 2007; Rodriguez-Caro et al. 2021). However, this analysis would require substantial further capture-mark-recapture data, ideally from multiple sites. Currently, we also only currently consider temporal variation within a homogeneous landscape, with all communities assumed to have the same demographic rates. Future work could test how spatial variation may impact on populations. In the UK, serotines are at the northern range-edge of their population. Understanding of how demographic parameters vary in space could allow for prediction of if, and how, this range edge will change under future climate scenarios. Spatial variation in demography may buffer some of the impacts of increasing environmental variation. If some colonies in high-quality areas have higher survival and reproduction, they may be able to produce dispersers that support lower-quality areas, leading to source-sink dynamics.

There remain a number of gaps in our understanding of serotine demography, as with many bat species, which limit the ability to realistically model these populations. The ranges assumed to be plausible for bat demographic rates were taken from analysis of a single roost. While the model was then calibrated to match to patterns expected for the wider population, it is possible that these original ranges were not representative of the wider population. Dispersal is expected to be a key driver of movement of both individuals and disease across populations. However, only limited information is available on bat movements, due to the challenge of tracking these small-bodied species. New, lighter tracking technologies may allow for greater elucidation of the movements of these species. A further gap is our understanding of density dependence in bat populations. Here, we assumed no density-dependent effects. However, Allee effects may be expected, given the benefits of larger maternity roosts for maintaining heat, or density-dependent effects on survival if there is resource competition, or increased parasitism within roosts. Inclusion of density dependence may alter the impacts of variation in demographic rates. We also assumed a fixed phenology, with processes such as birth and entry into hibernation pre-determined. In reality, bats show plasticity in their reproductive behaviour, including their ability to use torpor to delay parturition until conditions are suitable. Roost selection may also allow bats to move to locations which buffer environmental extremes.

The impact of inter-annual variation on the simulated serotine bat populations illustrates that estimates of mean survival and reproductive rates are insufficient to predict future population trends. Data from long-term studies which allow estimation of variation in these rates are therefore invaluable for identifying and predicting population trends. While serotine populations in the UK are currently stable, our results highlight that increasing environmental variation, particularly if it impacts adult survival, could lead to future population decline and influence disease persistence. Metapopulation dynamics appear to be central to persistence of this synthetic bat-borne zoonoses, emphasising the importance of understanding the movement dynamics of bat species for improved prediction for future disease prevention and control.

## Supporting information

S1 Model ODD

## Competing interests

The authors declare that they have no known competing interests.

## Acknowledgment

This work was funded by the Department for Environment, Food and Rural Affairs (Defra) under project code SE0432.

## Supporting information

Additional supporting information is provided as follows:

- **Appendix S1 (.pdf):** Model ODD. Full information about the model scope, design and operation including parameterisation, sensitivity analysis, and validation/verification.
- **Appendix S2 (.pdf):** Additional figures. Further illustrations of model output to support conclusions.

## Notes

### Competing Interest Statement

The authors have declared no competing interest.

