## Supplementary material for "Temporal variation in demography of temperate bats: consequences for population dynamics and disease": S1 Model ODD

### SimBat v1.0 - Serotine bat model ODD

#### 1. Purpose and patterns

This model simulates a population of Serotine bats (*Eptesicus serotinus),* and the transmission of a lyssavirus-like pathogen within this species. The model is structured into individual bat communities, each with a single maternity roost. Within each community, the survival, reproduction, and dispersal of individual bats is simulated, in addition to disease transmission. The purpose of the model is to assess the impacts of inter-annual environmental variation on bat population dynamics, and therefore the potential for disease spread.

The demographic model is calibrated by matching to the following patterns: a stable population size, the proportion of reproductive females in the maternity roost, and the population-level sex ratio.

#### 2. State variables and scales

The model simulates one type of agent, individual serotine bats. Bats are characterised by the following state variables:

**Demographic variables**

1) ID (Numeric ID)
2) community (Numeric ID)
3) sex (Male, Female)
4) age (Months)
5) age class (Pup, Juvenile or Adult)
6) reproductive status (Mature or Immature)
7) state (expired/ alive)
8) dispersal status (True, False)
9) maternity roost (True, False)
10) pups (list of pups as objects).

**Disease variables**

11) disease state (susceptible, latent, infectious, immune)

Bats are divided into communities, which occupy discrete spatial locations, represented by irregular polygons. Communities are characterised by:

1) ID (Numeric)
2) neighbours (adjacent communities)
3) bats_maternity_roost (list of individuals id within community occupying maternity roost).
4) bats_satellite_roost (list of individual ids within community occupying satellite roosts).

#### 3. Process overview and scheduling

Processes: Processes included in the model are reproduction, birth, survival, dispersal between communities and disease spread.

Scheduling: The model proceeds in monthly time steps across an annual cycle (Figure 1). Bat mortality and reproduction are expected to vary between years based on environmental conditions. Therefore, in each year, a relative quality was selected, depending on the scenario, which influenced reproductive and mortality parameters.

Following this, the model proceeded each year through each month in sequence (January to December). In April, bats are assumed to emerge from hibernation. At this point, juveniles become adults and females are tested for if they become reproductively mature (Reproduction). In June, birth is simulated, with pups produced based on fecundity (Birth). In all months, survival is then tested for all bats based on stage and season dependent parameters (Survival). In July, pups become independent juveniles. In October, maternity roosts are assumed to disperse, and all individuals are assumed to occupy smaller, satellite roosts. In October, natal dispersal is then simulated, allowing movement to adjacent communities (Dispersal). Disease processes are then simulated, with individuals progressing through pathology (latent, infectious, and immune period) and transmitting the disease (Disease). Finally, the list of bats in each community is updated to remove any bats which have expired and move dispersed bats before the next time step.

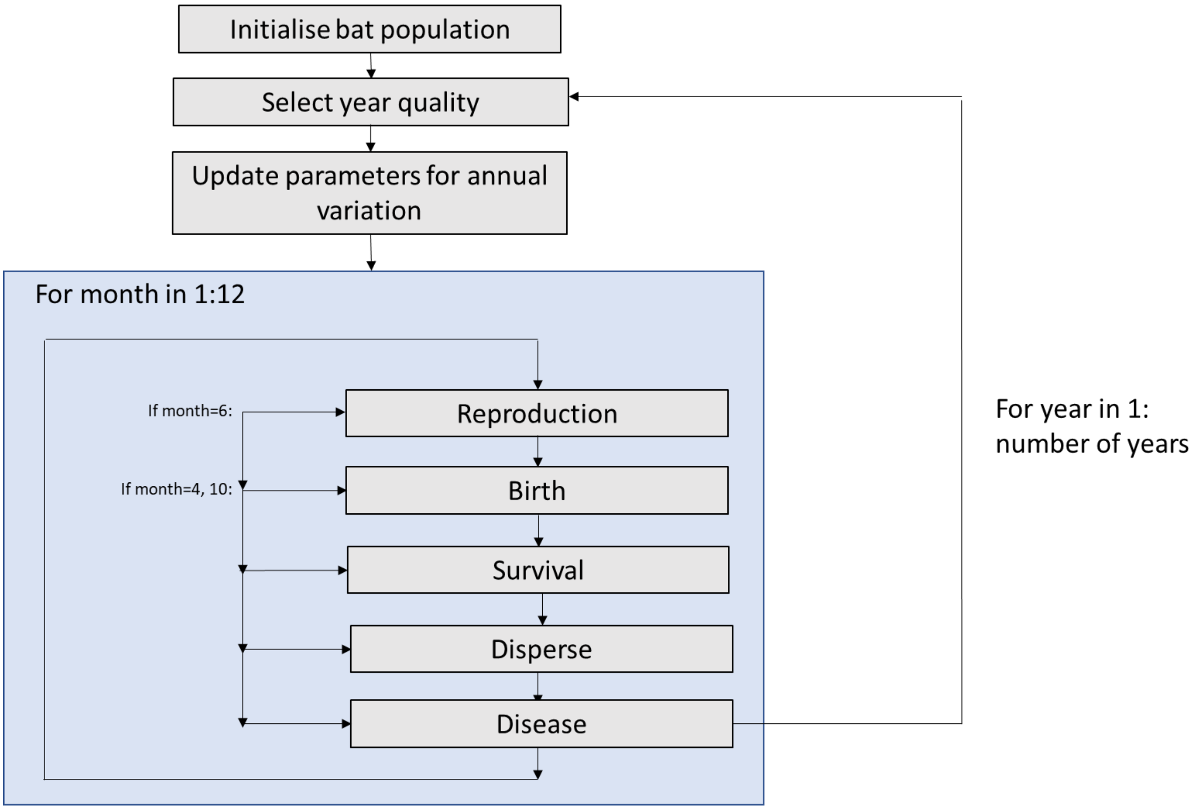


***Figure 1- Model diagram.***

#### 4. <u>Design concepts</u>

##### Basic principles

This model simulates individual Serotine bats, interacting within communities. The probability of reproduction, dispersal, mortality, and disease spread is stochastically determined at an individual level, allowing heterogeneity to emerge. Annual variation in mortality and reproductive parameters is also included.

**Emergence**

The population dynamics and age structure are an emergent property from individual reproduction, survival, and dispersal behaviours. Disease processes emerge from a combination of the population demography, dispersal and contact rates.

**Adaptation**

Not relevant

**Objectives**

Not relevant

**Learning**

Not relevant

**Prediction**

Not relevant

**Sensing**

Not relevant

**Interactions**

Disease spread results from interactions between susceptible and infectious bats.

**Stochasticity**

The majority of model processes were determined stochastically to represent environmental and demographic stochasticity. In each case, a random value between 0 and 1 was drawn, and events occurred if this value exceeded the probability assigned.

In addition, annual variation in mortality and reproduction was included based on a relative annual quality, which modified the default parameter.

**Collectives**

Bats are structured into communities, representing a collection of roosts shared by individuals which frequently interact. Breeding females, and a proportion of non-breeding females, were assumed to spend the active period in a single maternity roost per community, whereas males and other non-breeding females made use of satellite roosts. Roosts within communities were not explicitly simulated, but an individual-level variable was used to determine whether bats were in the maternity or satellite roosts, which was reflected in contact rates. Female bats are assumed to show high levels of philopatry, with most individuals remaining within these communities with limited dispersal between communities.

**Observation**

The total number of bats, and their age and sex distribution were tracked in each time step. The proportion present in maternity roosts relative to satellite roosts was also tracked. For disease processes, the number of infectious and immune individuals was tracked, and the proportion of communities in which disease was present.

#### 5. Initialisation

##### Community structure and carrying capacities

A total landscape of 30km by 30km was simulated. Within this landscape, bat communities were represented using contiguous irregular polygons (Figure 2). These landscapes were generated in R by randomly allocating a point per community (at a density of 0.05 per $km^2$, see data evaluation for justification) within a specified extent, and generating parcels by Voronoi tessellation. For each parcel, neighbours were identified, which determine potential inter-community transmission routes. A total of 10 different landscape randomisations were generated.

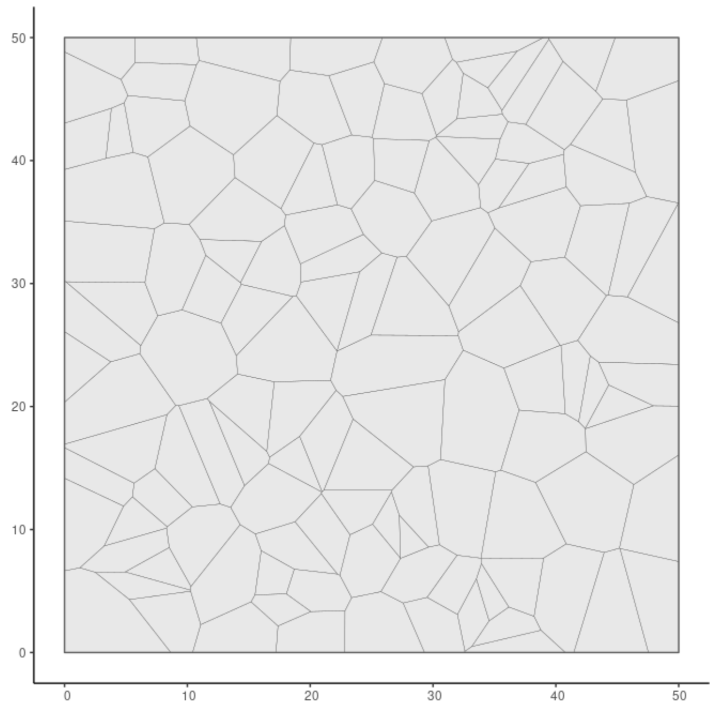


***Figure 2- Example model landscape- Polygons indicate community home ranges, each with a single maternity roost and a network of satellite roosts.***

The initial number of females in maternity roosts were randomly sampled from a negative binomial distribution based on BCT roost counts (see data evaluation). These populations were assigned to parcels so that the largest parcels had the largest initial population (Figure 3). At initiation, 50% of females were defined as reproductively mature. Males were added based on the sex ratio relative to the number of females for each community.

Each bat was assigned a sequential ID number. The model was run for a burn-in period of 20 years, in which results were not extracted, to allow demography to stabilise as age distributions emerge.

##### Disease seeding

Following a burn-in period, disease was seeded randomly across 20 bats. Infected bats were initially marked as being latently infected, with a latent period drawn from a Poisson distribution.

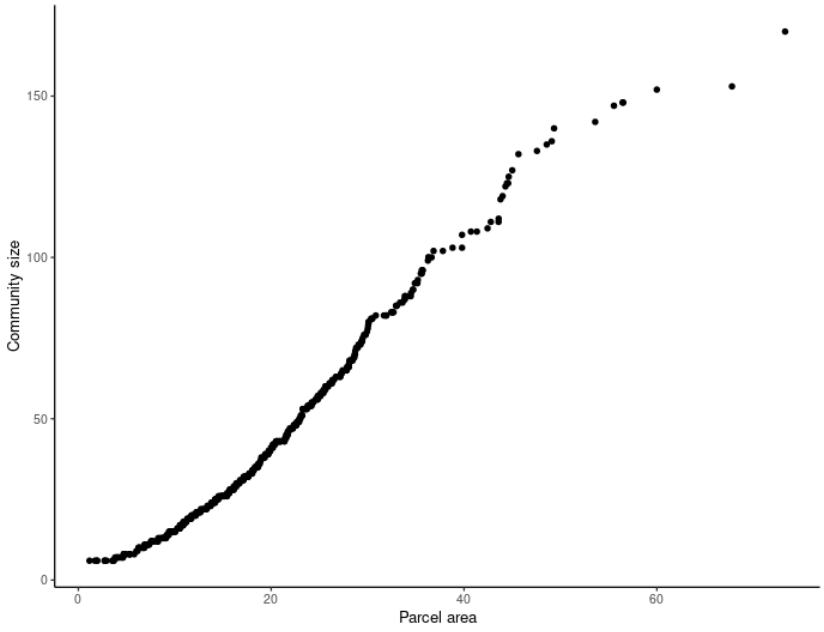


***Figure 3- Relationship between parcel area and allocated initial number of bats***

#### 6. Input data

##### Demographic

Plausible ranges for serotine demographic parameters were extracted from a Bayesian Capture-mark-recapture analysis from a single site across 25 years, in addition to the literature (Table 1). In some cases, parameters ranges were used for the closely related Big brown bat (*Eptesicus fuscus*).

***Table 1- Model parameters and ranges used in sensitivity analysis. Parameters in green were fixed, yellow were varied between specified ranges.***

| Parameter | Value | Uncertainty range | Unit | Source |
|---|---|---|---|---|
| **Community seeding density** | 0.05 | 0.004-0.12 | Communities per $km^2$ | Median value estimated by Mathews et al. (2018). |
| **Total number of communities** | 45 | NA | | Based on density of 0.05 communities per km2 across 50km-by-50km arena. |
| **Community sizes** | Negative binomial distribution (μ=41.2, size=1.65) | Fitted to data | Roost carrying capacity | BCT maximum roost counts. |
| **Hibernation period** | October-March (10-3) | NA | Month | Period during which bats assumed to be in torpor, resulting in lower mortality |
| **Active period** | April-September (4-9) | NA | Month | Period during which bats active, and therefore vulnerable to higher mortality |
| **Breeding period** | June-July (6-7) | NA | Month | Period during which assumed higher mortality |

| | | | | |
|---|---|---|---|---|
| | | | | for breeders due to stress of reproduction |
| **Birth month** | June (6) | NA | Month | |
| **Dispersal months** | October (10). | NA | Months | Assumption that juvenile dispersal will occur pre- and post-first hibernation, as juveniles relocate with other individuals from same hibernacula. |
| **Sex ratio at birth** | 0.5 | Not considered | Probability that pup is male | Assumption of equality. *E.fuscus* sex ratio at birth approximately even but varies with time of reproduction (Barclay 2012) |
| **Female dispersal probability.** | 0.02 | Not considered | Per-year probability that reproductively immature female bat disperses | Assumption based on evidence for natal philopatry |
| **Juvenile primiparity** | 0.12 | 0.05-0.26 | Per year probability that female becomes reproductively mature as a juvenile (age= 1 year). In subsequent years, multiplied by age to give probability. | CMR analysis of Crundale site<br>Harbusch (2003) reported that serotines reach reproductive maturity by the end of their second year, but under favourable conditions may in first year. |
| **Non-breeder primiparity** | 0.24 | 0.20-0.39 | Per year probability that female becomes reproductively mature as a previous non-breeder (age> 1 year). In subsequent years, multiplied by age to give probability. | CMR analysis of Crundale site |
| **Fecundity** | 0.93 | 0.84-0.97 | Per year probability that reproductively mature female produces pups | CMR analysis of Crundale site<br>In *E. fuscus*, O'Shea et al. (2010) found that given previous breeding, *E. fuscus* had a 0.96 (95% CI: 0.94-0.98) conditional probability of reproducing the next year. |
| **Twinning** | 0.02 | 0-0.05 | Per-female probability of second pup being produced | Assumption- twins rarely observed. |
| **Pup survival** | 0.94 | 0.7-1 | Probability that dependent pup survives first month to weaning | Harbusch and Racey (2006)- 0-27% mortality of pups per year. |
| **Juvenile survival** | 0.53 | 0.48-0.83 | Probability of survival to recapture in year 1. Converted to per month probability. | CMR analysis of Crundale site<br>In *E.fuscus* O'Shea et al. (2010) estimated first year survival as 0.67 (95% |

| | | | | |
|---|---|---|---|---|
| | | | | CI:0.61-0.73) for weaned females. |
| **Non-breeder survival** | 0.91 | 0.69-0.93 | Annual survival probability. Converted to constant per month probability. | CMR analysis of Crundale site (Robardet et al. 2017) Site A- 0.86 [0.76–0.93]. Site B- 0.78 [0.73–0.83] In *E.fuscus* O'Shea et al. (2010) estimated adult survival as 0.79 (95% CI:0.77-0.81) for weaned females. |
| **Breeder survival** | 0.83 | 0.70-0.90 | Annual survival probability. Converted to per-month survival penalty during reproductive period (April-August) for reproductively mature females. | CMR analysis of Crundale site |
| **Proportion of non-breeders at maternity roost** | 0.68 | 0.1-0.9 | Probability that non-breeding female uses maternity roost | Assumption |

**Seasonal dynamics**

Bats were assumed to enter hibernation in October, and emerge in April (BCT serotine fact sheet; Robinson and Stebbings 1997). Following hibernation, breeding females were assumed to return to maternity roosts for the period from April-August, where they form large colonies. Females defined as reproductively mature remained solely in the maternity roost through the reproductive period (June-July), whereas a proportion of non-reproductive females used satellite roosts (Catto et al. 1996). Males were assumed to be present in the same communities, but not utilising the primary maternity roost, instead using satellite roosts throughout the active season (Table 2). In the autumn pre-hibernation period (September), maternity roosts were assumed to break up, and all individuals to roost in satellite roosts throughout the community.

Pups are assumed to all be born in June. A study of serotine maternity colonies in Germany found a mean birth date of 16$^{th}$ June, with most births occurring within a two-week (14±6 days) period (Harbsuch and Racey 2006). Catto (1993) found pups were born between the 29$^{th}$ June and 6$^{th}$ July on the south coast of England. Pups were assumed to be dependent for a single month and became independent juveniles in July. The pup class therefore represents the period within which the bat is largely confined to the single maternity roost, and spans birth to first emergence. Whilst we understand that there may be a small overlap between the first flights and full weaning and independence, we assume here that this is less than one month and is generally assumed to be best counted as a few days, rather than in weeks. Thus, for simplicity we assume first flight and independence are simultaneous. Harbusch and Racey (2006) found that young bats first emerged 36±7 days after birth. Given a mean birth date of mid-June, this supports emergence of juveniles in July.

The juvenile class is defined as starting with independence and ending at the end of first hibernation. Following first hibernation, it is unlikely that the morphology of an unmarked bat would permit its description as the young of the previous year.

Dispersal of juveniles between communities was assumed to occur during September, prior to hibernation. During hibernation, it was assumed that no contact between bats occurred, and that pathology processes were paused.

*Table 2- Presumed location of bats by class across year*

| Month | Breeding Female | Non-breeding Female | | Juvenile M/F | Adult Male | Month specific processes |
|---|---|---|---|---|---|---|
| Jan | Hibernating | Hibernating | | Hibernating | Hibernating | **Hibernation** |
| Feb | Hibernating | Hibernating | | Hibernating | Hibernating | **Hibernation** |
| Mar | Hibernating | Hibernating | | Hibernating | Hibernating | **Hibernation** |
| Apr | Maternity roost | Maternity roost | Satellite roost | | Satellite roost | |
| May | Maternity roost | Maternity roost | Satellite roost | | Satellite roost | |
| Jun | Maternity roost | Maternity roost | Satellite roost | | Satellite roost | **Pups born** |
| July | Maternity roost | Maternity roost | Satellite roost | Maternity roost | Satellite roost | **Pups become independent** |
| Aug | Maternity roost | Maternity roost | Satellite roost | Maternity roost | Satellite roost | |
| Sep | Satellite roost | Satellite roost | Satellite roost | Satellite roost | Satellite roost | **Breeding/move to hibernation sites** |
| Oct | Hibernating | Hibernating | | Hibernating | Hibernating | **Dispersal of males and non-breeding females, Hibernation** |
| Nov | Hibernating | Hibernating | | Hibernating | Hibernating | **Hibernation** |
| Dec | Hibernating | Hibernating | | Hibernating | Hibernating | **Hibernation** |

##### Community distribution

Serotine bats are limited in distribution to the south of England and Wales and primarily rely on roof spaces for maternity roosts. Mathews et al. (2018) reviewed different estimated densities for maternity roosts and found a median of 0.05 roosts/km$^2$ within the species range, with a range of 0.004-0.12 roosts/km$^2$.

Using a seeding density of 0.05 roosts/km$^2$ gives a mean area for each community of 20km$^2$. This estimate broadly aligns with empirical estimates for bat home ranges. Figure 4 shows the distribution of community sizes from the model relative to a distribution of home range estimates from serotine bats from a global database of mammalian home ranges (Broekman et al. 2022). While the footprint of communities will differ from that of individual bats, this suggests the spatial scale of the modelled landscape is appropriate.

##### Maternity roost size

Initial maternity roost sizes allocated to each parcel were based on maximum roost counts from the BCT. The counts are carried out in June, prior to juveniles leaving the roost, and therefore represent the maximum number of adults observed. For serotines across all years, the mean count was 36 and median 24. It was assumed that very small roost counts (<5 individuals) did not represent an entire maternity roost, therefore these counts were excluded from the data. A negative binomial (μ=41.2 and size=1.65) was fitted to the data using the fitdistr package in R (Figure 2). Community sizes were allocated to parcels in order of parcel area (largest parcels had largest maternity roost size). Across 10 landscapes of 500 communities, this gave a mean bat density of 1.94 individuals/km$^2$ for individuals in maternity roosts pre-breeding. Assuming this is doubled to account for males (3.88 individuals/km$^2$), this falls within the plausible range for individual bat density from Mathews et al. 2018 (0.1-4.6 bats/km$^2$).

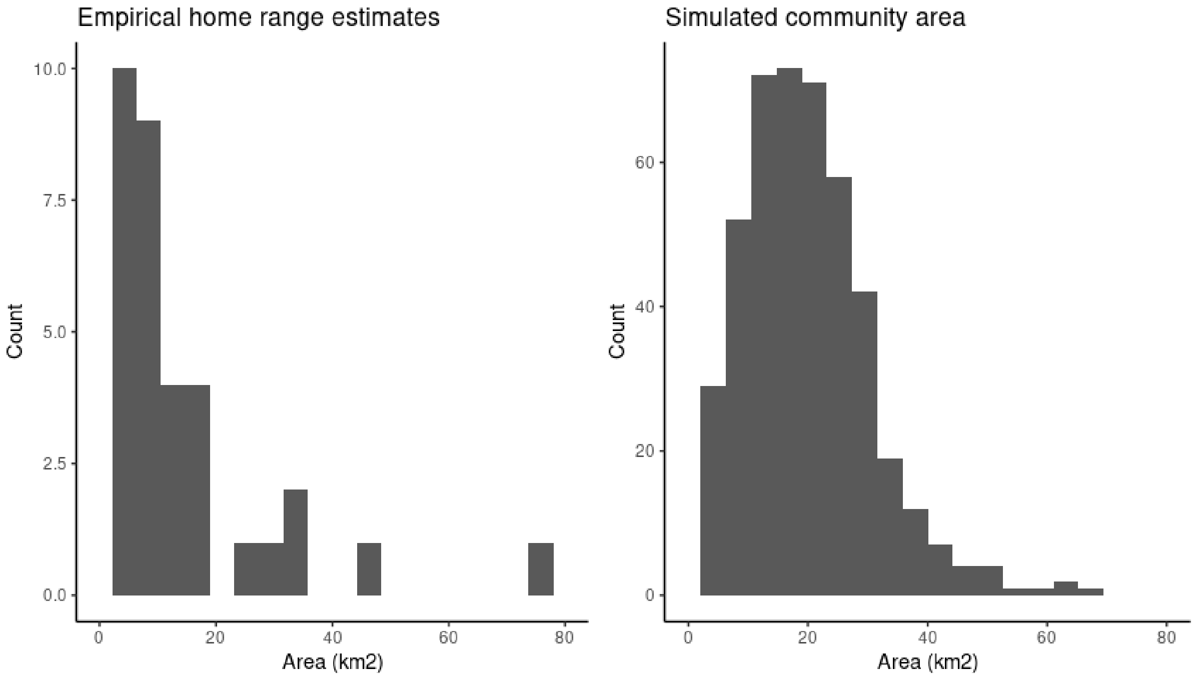


***Figure 4- Distribution of serotine home range areas from global database (Broekman et al. 2022) relative to simulated community areas across 10 landscapes.***

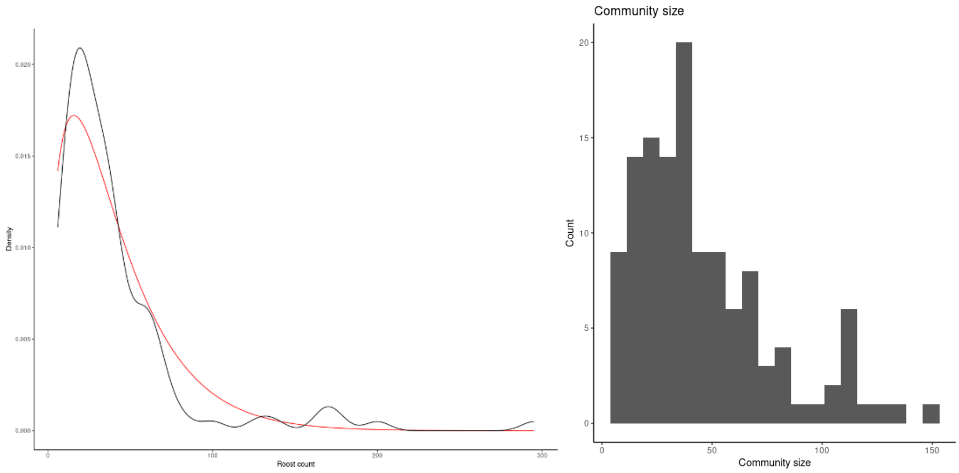


***Figure 5- A) BCT serotine maximum roost counts. Black line shows density from data set and red line indicates fitted negative binomial distribution. B) Distribution of community sizes for single model simulation.***

#### Primiparity

The primiparity parameter defines the per-year probability of a bat becoming reproductively mature. Two separate probabilities were used: the probability that a juvenile reproduces in its first year (primiparity_juv) and the probability that an adult non-breeder (age>1 year) becomes a breeder (primiparity_nb).

Chauvenet et al. (2006) found a long pre-reproductive period in female serotines, with a median duration of 3.5 years (maximum of 14 at Hollingbury and 10 at Crundale) between individuals being caught as juveniles and being observed to breed. However, these estimates may be biased by recapture rates as bats were only caught at a maternity roost, which is more likely to be used by breeders than non-breeders. Harbusch (2003) reported that serotines reach reproductive maturity by the end of their second year, but under favourable conditions may in first year. Similar trends have been reported in *E.Fuscus,* with females born earlier the previous year, more likely to reproduce as a one-year-old (Barclay 2012).

Plausible ranges for the primiparity parameters were extracted from a Bayesian analysis of a capture-mark-recapture dataset of a roost in Crundale, Kent (Table 1).

**Fecundity**

The fecundity parameter defines the per-year probability of producing a pup for reproductively mature females. CMR analysis of the Crundale site estimated a fecundity (defined as remaining in the breeding class) of 0.92-0.996. In the closely related *E. fuscus*, O'Shea et al. (2010) found that given previous breeding, *E. fuscus* had a 0.96 (95% CI: 0.94-0.98) conditional probability of reproducing the next year, within the range estimated for *E.serotinus.* We assumed that fecundity was independent of roost size, however in *E.Fuscus* Mills et al. (1975) found evidence for density dependence, with lower fecundity as roost sizes increased.

**Twinning**

Twins have been observed in serotines, but relatively rarely (Aegerter, pers. comm), therefore a plausible range of 0-5% of females producing twins was assumed.

**Sex ratio**

Sex ratio at birth was assumed to approximate equality. Catto (1993) reported no significant difference from 1:1 in a sample of 87 captured juveniles. In *E.Fuscus,* Barclay (2012) reported variation in sex ratio within years, with female-biased births earlier in the season in years where parturition occurred early. However, the overall sex ratio did not differ significantly from 1.

**Dispersal and carrying capacities**

Limited information is available on dispersal of serotines between communities due to the difficulties in directly observing bat movements. In general, temperate bats are considered to show high female natal philopatry, with gene flow driven by male inter-colony movements or mating. This is also supported in serotines, on the basis of sex-specific markers (Moussy et al. 2015). Levels of gene flow between populations appear to be relatively high, although lower in the UK than for continental European populations (Moussy et al. 2015). In Poland, in a genetic analysis only 22% of individuals were assigned to the sub-population from which they were sampled. Individuals defined as migrants had ranged distances from 60-283km in females, and 27-385km in males (Bogdanowicz et al., 2013).

In the absence of more detailed information, we made the following assumptions:

- All males, in the October of their first year (prior to first hibernation) randomly select a community to move to from across the arena.
- Once females become reproductively mature, they remain within their current community. However, reproductively immature females have a low (0.02) probability of moving

community each October. As with males, they randomly choose a community to move to from across the arena.

**Movement between maternity and satellite roosts**

During the reproductive period (May-August), maternity roosts consist solely of females and their offspring from that year. Reproductive females require warmer roosts to reduce use of torpor and promote foetal development. It was assumed that all breeding individuals will use a single maternity roost within the community throughout the active period. By contrast, non-breeding females, which have different energy requirements, may choose to use either the maternity roost or alternative 'satellite roosts' within the community. This was defined stochastically for each individual on emergence from hibernation in April. Little is known about the movements of male serotines, however, they are also likely to use satellite roosts within communities, roosting individually or in small groups. Maternity roosts typically disperse in August. After this point it was assumed that all bats within the community are making use of networks of smaller roosts throughout the community.

Relatively little is known about mating behaviour in serotines. However, observation of a roost in a church in the Netherlands observed mating starting in September, after weaning of pups, and peaking in October, with last observations in December. Some mating was also observed in March and beginning of April (Fasal at al. 2023).

**Pup survival**

Pups were assumed to all be born in June, and to be dependent on mothers for a single month prior to fledging in July. Prior to weaning, if their parent dies, pups were also assumed to expire.

Harbusch and Racey (2006) found that pup mortality primarily occurred in young pups (<10 days) and during periods of inclement weather, with 11 to 27% of pups dying. In a year with no periods of poor weather, mortality was 0%. Glas (1981) reported mortality of 21-29% of juvenile serotine bats. These values will include both mortality due to the death of the mother, and survival of the mother but death of the pup due to other factors. A broad possible range of pup mortality from 0-30% was therefore considered, which determines survival of a pup, dependent on maternal survival.

**Juveniles and adult survival**

Survival of bats was tested in each monthly time-step. For annual estimates, plausible parameter ranges were based on the CMR analysis from the Crundale roost. Survival was estimated separately for juveniles, non-breeders/unknown and breeders from one year to the next, with survival found to be lower for juveniles and breeding individuals. No estimates were available for survival of adult males, therefore in the absence of additional information, adult males were assumed to be equivalent to non-breeding females, on the assumption that males do not face the high energy requirements of breeding females.

Juvenile and adult survival is expected to vary within years, due to seasonal variation in behaviour and environmental conditions. Mortality is expected to be lower during hibernation than the summer period, due to reduced risks of predation (Turbill and Ruf 2011; Reusch et al. 2019) and lower energy requirements. Survival is also expected to be reduced for breeding individuals due to the energetic costs of pregnancy and lactation. To account for this variation, annual survival probabilities were converted to monthly probabilities based on the following:

##### *Adults*

To account for seasonal variation, using a given annual adult survival probability (AS), mortality was divided into a number of 'shares', where:

$$AS = M^{shares}$$

If mortality is equally probable in all months, shares is equal to 12, therefore monthly mortality (M) is equal to $AS^{(1/12)}$.

We implemented seasonal variation in mortality by allocating more 'shares' of mortality to summer months. On the assumption that the summer season is 6 months (April-September) and bats hibernate (October-March), for 6 months:

$$AS = SS^{6*share\_increase} HS^{6}$$

Where SS is summer monthly survival, HS is hibernation monthly survival and share increase is the number of 'shares' of mortality allocated during the summer relative to hibernation. This value can be interpreted as the number of months in hibernation it would take to equal the mortality risk of a single summer month.

In *E.Fuscus*, O'Shea et al. (2011) estimated a weekly survival of 0.998 in winter, and 0.984 during summer, equivalent to a monthly survival in summer of 0.933 and in winter of 0.991. For this species, If HS is therefore taken as 0.991, the share increase in summer can be calculated based on:

$$Summer\_share\_increase = \frac{\ln(\mathrm{SS})}{\ln(HS)}$$

Therefore, giving an estimate of 8.05. A summer share increase of 8 was therefore used. For simplicity, we assumed this relative rate of summer to winter mortality remained constant. In Natterers' and Daubenton's bats, Reusch et al. (2019) found a consistent pattern between seasons that varied between years, as modelled here.

Therefore, in summer months, survival is equal to:

$$SS = (AS^{1/num\_shares})^{Summer\_share\_increase}$$

And during the months of hibernation:

$$HS = (AS^{1/num\_shares})$$

As an example, given an annual mortality of 0.905, the summer monthly mortality is 0.985 and winter mortality is 0.991.

##### *Juveniles*

Bats are defined as juveniles from when they leave the roost in July until they emerge from hibernation the following April. The same logic as above was used to convert annual survival probabilities for juveniles to seasonally dependent monthly probabilities, however juveniles are only defined as such for 9 months (3 summer and 6 during hibernation). The following equation can therefore be used to estimate summer and winter juvenile mortality:

$$JS = SS^{3*share\_increase} HS^{6}$$

From April, juveniles are not separable from adults, therefore adult survival rates were applied.

***Breeders***

Survival for breeding individuals was estimated to be lower than for non-reproductive individuals. This was assumed to be due to the stress relating to late-pregnancy and lactation, and was therefore implemented as an increase in mortality during the maternity period (June-July), with mortality equal to non-reproductive individuals outside of this period (4 remaining months of summer and 6 months of winter). Therefore, for breeders their annual mortality (AS) is equal to:

$$AS = SS^4 HS^6 BS^2$$

Where BS is the monthly survival during breeding. The survival during those breeding months (BS) is therefore calculated as:

$$BS = \frac{AS}{SS^4 HS^6}^{\frac{1}{2}}$$

Figure 8 shows the cumulative survival probability across a year for a juvenile, non-reproductive adult, and breeding individual, including seasonal variation.

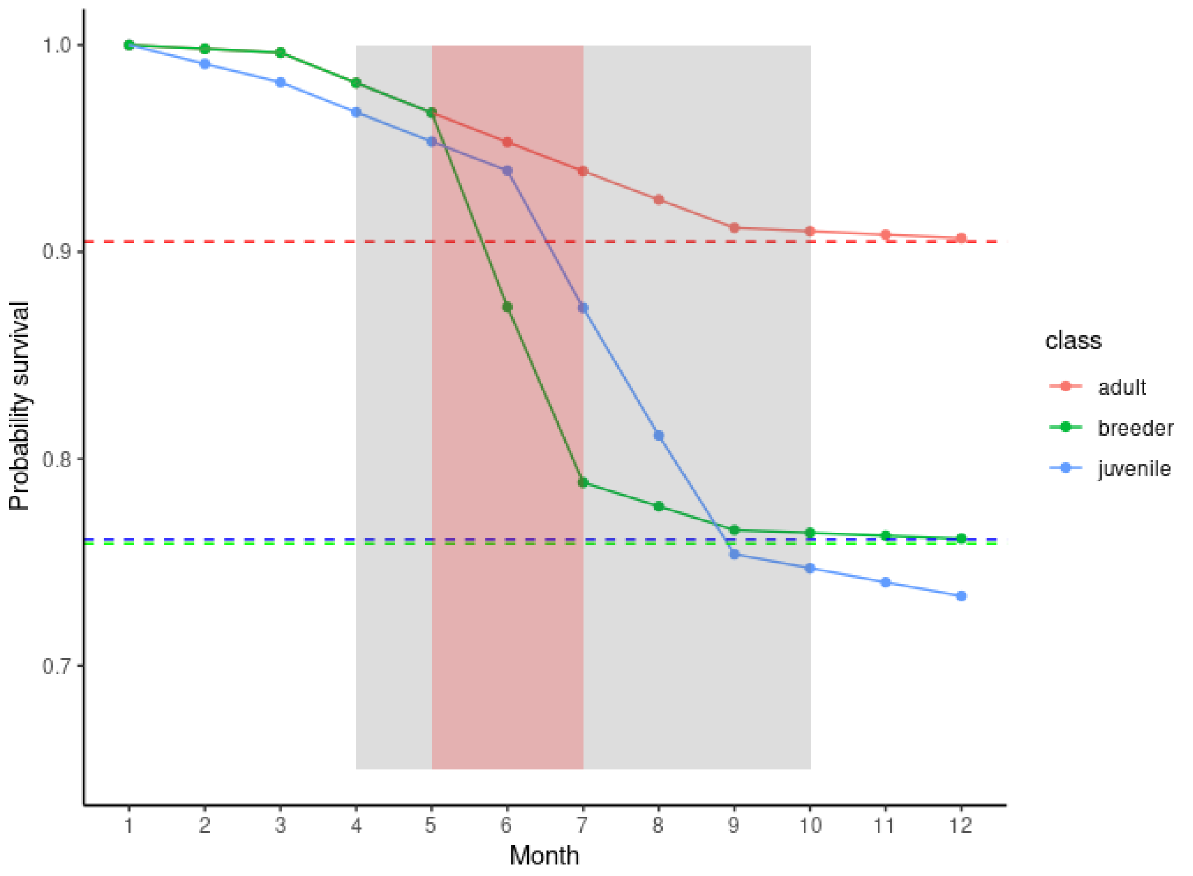


***Figure 6- Seasonal variation in mortality for different classes. Red indicates the breeding window, and grey the active summer season.***

#### <u>Disease input data</u>

Disease parameters were chosen based on literature estimates, both for EBLV-1 in serotines and for lyssaviruses in other bat species.

| Parameter | Value | Uncertainty | Unit | Source |
|---|---|---|---|---|
| Initial number exposed | 20 | Model assumption | Number of bats initially latently infected | Assumed |
| Transmission probability | 0.75 | | Probability of disease transmission per contact | Assumption to give stable dynamics |
| Probability of becoming infectious | 0.15 | | Probability that infection leads to mortality | Consistent with estimates for lyssaviruses in other bat species.<br>0.1<br>(RABV-*Desmodus rotundus* Blackwood et al. 2013)<br>0.15<br>(RABV- *Eptesicus fuscus* George et al. 2011)<br>0.058<br>(EBLV-1 *Myotis myotis*) |
| Duration latency | 1 | | λ for Poisson distribution for latent period | Consistent with assumed mean rates of 21-24 days for bat-lyssavirus models (Blackwood et al. 2013; Kim et al. 2023; George et al. 2011) |
| Duration infectious period | 1 | | Months. Assumed infectious period considerably shorter than month time step. | Consistent with assumed mean rates 5-6 days for bat-lyssavirus models (Blackwood et al. 2013; Kim et al. 2023; George et al. 2011; Amengual et al. 2007) |
| Duration immunity | 6 | | λ for Poisson distribution for immunity period | Assumed. Estimates and assumptions for bat-lyssavirus systems are highly variable, ranging from ~4.5 months (Blackwood et al. 2013) to greater than a year (Amengual et al. 2007; Colombi et al. 2019) |
| Contact rate (maternity roost) | 2.5 | | Contact rate for individual in maternity roost per month | Assumption on the basis of high rates of contact within maternity roosts. |
| Contact rate (satellite roosts) | 0.25 | | Contact rate for individual in satellite roost per month | Assumption on the basis of low rates of contact within maternity roosts. |
| Contact rate (neighbours) | 0.05 | | Contact rate between individuals in satellite roosts in neighbouring communities | Assumption on the basis of low rates of contact between neighbouring communities. |

**Probability of developing clinical infection**

Experimental lyssavirus infections in bats, as well as field studies, have suggested the potential for different pathologies. Potential courses of infection include:

- Classical- exposure leads to latent period followed by development of clinical infection which is inevitably lethal (SEI)
- Abortive/subclinical- infection leads to development of an immune response and clearance of the virus without becoming infectious (SER)
- Recovery- infection leads to clinical infection, followed by clearance and recovery (SEIR)

- Intermittent infection- Infection leads to cycles of latency and infectivity
- Carrier state- Infection leads to extended period of infectiousness, without symptoms

For simplicity, we assumed only classical and abortive infection occurred. Following exposure, bats entered a latent period. At the end of this period, bats either become infectious, and inevitably die, or clear infection and become immune. We assumed 15% of infections would lead to clinical infection, which is comparable with estimates for RABV in *Demodus rotundus* (~0.1, Blackwood et al. 2013), RABV in *Eptesicus fuscus* (0.15, George et al. 2011) and EBLV-1 in *Myotis myotis* (0.058, Kim et al. 2023).

**Latent period**

Latent periods for rabies can be highly variable. Previous modelling studies have used a mean latent period of less than a month (24 days- George et al. 2011; 24 days- Kim et al. 2023; 21 days- Blackwood et al. 2013). However, latent periods are extended by the use of torpor (Davis et al. 2016). Latent periods were drawn from a Poisson distribution with a mean of 1 month. We assumed that the latent period only progressed during the active summer period, based on evidence from experimental infection of other lyssaviruses (Davis et al. 2013).

**Infectious period**

Infectious periods for rabies tend to be short. Previous modelling studies for lyssaviruses in bats have assumed infectious periods of less than a week (6 days- George et al. 2011; 5 days- Kim et al. 2023; 5.78 days- Blackwood et al. 2013; 5.1 days- Amengual et al. 2007). On this basis, we assumed a bat would only remain infectious for a single time step, following which it would succumb to the disease.

**Duration immunity**

Duration of immunity in bats is challenging to estimate as it requires sequential sampling of the same individuals. For RABV in *Desmodus rotundus,* Blackwood et al. (2013) assumed a mean of 4.5 months to lose immunity, whereas for *Myotis myotis* and *Myotis schreibersii*, Colombi et al. (2019) predicted mean immunity greater than a year. Persistence of EBLV-1 seropositivity for more than a year has also been found in a field study of *Myotis myotis* (Amengual et al. 2007). Kim et al. (2023) found that models where immunity was lost at a faster rate over hibernation provided a better fit to data for *Myotis myotis,* with 82% of bats losing immunity during hibernation, relative to 8.4% before hibernation.

We assumed a mean duration of 6 months, with the duration for each individual drawn from a Poisson distribution, allowing longer durations to also occur.

**Contact rates and transmission probabilities**

We assumed contact rates would vary depending on whether individuals were present in maternity roosts, or satellite roosts. Maternity roosts host large numbers of individuals in close contact. Relatively little information is available on non-maternity roosts, used by males, non-breeders, and breeding females outside of the active period. However, given that high temperatures are not required by these individuals, it was assumed individuals are roosting either singly, or in small groups. It was also assumed that bats in satellite roosts may also contact individuals from neighbouring communities at a lower probability.

Contact rates were assumed to be intermediate between density and frequency dependent so contact rates increased with community sizes (in maternity roosts or satellite roosts independently) but reached a plateau. Contact rates will vary seasonally based on the number of individuals in maternity roosts and satellite roosts.

#### 7. <u>Sub-models</u>

##### Initialise

This function generates the initial bat population and community structure. A list of communities is generated, with each community represented as an object and assigned a list of neighbouring communities to which disease can be transmitted.

A list of bats is the generated, with all bats initially 36 months old, and a sex ratio of 40:60 females to males. 50% of females are initiated as reproductively mature.

##### Convert survival parameters

This function takes the input annual survival parameters (as logits) and converts them to monthly estimates based on the class, and year quality.

##### Reproduction

The function determines the development of reproductive maturity, and movement into and out of the maternity roost.

In April (Month 4), juveniles are defined as adults. For females that had not previously reproduced, whether they became reproductively mature is then stochastically determined based on the primiparity parameter, defined separately for juvenile females and previous non-breeders. If this condition was met, bats were defined as reproductively mature and remained in this state. All reproductively mature females at this point are assumed to move into the maternity roost for the summer period. For immature females, whether they choose to move to the maternity roost, or use satellite roosts, is stochastically determined.

In September (Month 9), all bats are assumed to all leave the maternity roost for the year.

##### Birth

This function is called only in June (Month 6). The functions loops through all bats, and for each reproductively mature female, stochastically determines whether they successfully produce a pup based on the fecundity parameter. This fecundity parameter is calculated for each year based on the year 'quality'. Serotines are promiscuous, and it was therefore assumed that mating was not a limiting factor. For each female that reproduced, there was a probability of twinning. For each pup produced, sex was stochastically determined, and a sequential ID number was assigned. Pups were added as objects to the female. The function returns the updated list of bats with pups included, the current id number, the total number of females reproducing, and the total number of pups produced.

##### Survival and ageing

Each month, all bats are aged by 1. For reproductively mature females, if they have pups, these are also aged, and once they reach two months old (July), become independent juveniles, which are removed as objects from the litter list and added to the primary bat list.

Survival is then determined stochastically, based on a probability specific to each particular age class (pup, juvenile, adult). For reproductively mature females, survival was also dependent on whether they are breeding, with higher mortality assumed to occur due to the stress of reproduction during the breeding period (June to July). If a female with pups dies, her pups were also marked as expired.

The function returns the updated list of bats, the ages of dead bats and the number of dead bats of each class.

**Disperse**

This function is called in October (Month 10 only). All bats marked as males and juveniles (Age <1 year) were assigned to choose a random community to settle in. For bats marked as females and reproductively immature, it was assumed there was a low probability of dispersal with whether each individual dispersed stochastically determined. If dispersal occurred, a random community was chosen to move to.

**Update communities**

This function updates the list of bats in each community and counts the number which are in both the maternity roosts and satellite roosts and the number of infected.

**Seed infection**

This function is called after the burn-in period to seed disease. A user-specified number of bats are randomly sampled from across communities and assigned as latently infected, with a latent period drawn from a Poisson distribution.

**Transmission**

A generic endemic disease was simulated within the bat population. Pathology was simulated as a SEIR process (Susceptible, Exposed/Latent, Infectious, Recovered/Immune).

For each bat, whether they are infectious is first tested. Following 1 month as an infectious, these bats are marked as expired.

For bats which are latently infected, a countdown is used to measure the time to either becoming infectious or immune. This countdown only progresses during the active period. When this countdown reaches 0, whether bats become immune or infectious is stochastically determined based on the Probability of becoming infectious parameter. For bats which become immune, a duration of immunity is drawn from a Poisson distribution. This duration is used as a countdown timer. When this reaches 0, bats are marked as ‘susceptible’ and can become re-infected.

For bats which become infectious, in this model version, the infectious period is assumed to be short-lived relative to the time-step. At the end of the time-step the number of infectious bats is counted within each community, and this is used to define the force of infection in the next time step.

The disease was assumed to be directly transmitted, with a transmission function chosen to be intermediate between density and frequency dependent. We assumed three different contact rates depending on the likelihood of contact between different groups:

- Maternity roost: contact rate for individuals within maternity roosts (juveniles, breeding females, and a proportion of non-breeding females) during summer period.
- Satellite roosts: contact rate for individuals within a community using satellite roosts

- Between-community: contact rate for individuals using satellite roosts with individuals from neighbouring communities' roosts

The following equation was used to determine the probability of a susceptible individual becoming exposed (p_inf) within a single time step, for individuals within the maternity roost:

$$p(\text{inf}) = 1 - (1 - p_{trans})^{C_{MR} I_{MR} / N_{MR}^{0.5}})$$

Where $C_{MR}$ is a mean contact rate between individuals within maternity roosts, $I_{MR}$ is the number of infected individuals within the maternity roost, $N_{MR}$ is the total number of individuals within the maternity roost and $p_{trans}$ is the probability that transmission occurs for a given contact.

For individuals in satellite roosts, the following was used:

$$p(\text{inf}) = 1 - ((1 - p_{trans})^{C_{SR} I_{SR} / N_{SR}^{0.5}} \prod_{j=0}^{n} (1 - p_{trans})^{C_N I_N / N_N^{0.5}}$$

Where $C_{SR}$ is a mean contact rate between individuals within satellite roosts within a community, $I_{SR}$ is the number of infected individuals within the satellite roosts, $N_{SR}$ is the total number of individuals within the satellite roosts, $C_N$ is a mean contact rate with individuals in neighbouring satellite roosts, $I_N$ is the number of infected individuals within neighbouring satellite roosts and N is the total number of individuals within neighbouring satellite roosts.

For each individual, whether it became exposed was stochastically determined based on the maternity roost, or satellite-roost probability. Exposed bats are assigned a latent period drawn from a Poisson distribution.

At the end of the simulation, infectious bats which are marked as expired are removed from the list of bats, so they are not re-counted for the force of infection in the next time-step.

#### 8. Implementation

##### Software

This model was implemented in python version 3.8.10.

#### 9. Pattern orientated matching

The following patterns were used for model calibration and validation

##### Population size change

Population stability was matched against information from the Bat Conservation Trust (BCT) National Bat Monitoring Programme (NBMP), which suggests serotine populations are relatively stable. Roost counts suggest a mean annual decrease of 0.03%, but with a confidence interval encompassing 0 (95% CI: -1.5% to 1.8%.).

##### Sex ratio

The majority of studies of serotines have been carried out at maternity roosts, where adult males are excluded. To estimate the population-level sex ratio, we used published estimates from passive surveillance schemes for disease testing of bats (Table 3). These submissions should represent relatively random samples across serotine populations. This method was used by Kurta and Matson (1980) to estimate sex ratio for *E. Fuscus*, with a male-biased sex ration of 61:39 found (220:142 males to females). Results from serotine studies support that serotine populations are also male biased, with across different studies, a sex ratio of 60:40 males to females (928 males to 623 females).

**Proportion reproductive**

For the proportion of breeding females, relatively limited information is available. In a study at maternity roosts, Catto et al. (1996) reported that of adult females captured, 66% were reproductively active. Based on an available dataset of serotine captures within the UK (914 annual captures of adult females with 682 unique individuals- repeated captures within a year were excluded), 614 captures were defined as reproductively active, 161 as non-reproductive and 139 as unknown. Depending on the status of the unknown class, this gives a proportion of reproductive individuals ranging from 67-82%. Assuming un-defined individuals are more likely to be non-reproductive, and this estimate aligns closely with that from Catto et al. (1996), we assumed 66% of females within maternity roosts were reproductive as a best estimate.

***Table 3- Studies reporting sex ratios from passive surveillance schemes***

| **Source** | **Number of males** | **Number of females** | **Sex ratio (M:F)** | **Location and method** |
|---|---|---|---|---|
| Van der Poel et al. (2005) | 706 | 473 | 60:40 | Passive surveillance from Netherlands 1984-2003. Both positive and negative included. |
| Harris et al. (2006) | Not reported. Total of 82 bats so presume 46 based on ratio. | Not reported. Total of 82 bats so presume 36 based on ratio. | 56:44 | Passive surveillance from UK 1987-2004. Both positive and negative included. |
| Muhldorfer et al. (2011) | 21 | 13 | 62:38 | Passive surveillance from Germany 2002-2009. Both positive and negative included. |
| Picard-Meyer et al. (2016) | 92 based on 21 positive males, reported as 22.9% of submitted | 77 based on 16 positives, reported as 20.8% of submitted | 54:46 | Passive surveillance from France 1989-2013. Both positive and negative included. |
| APHA Bat Rabies Dashboard | 39 | 25 | 61:39 | Passive surveillance from UK 2019-2022. Both positive and negative included. |
| Orlowska et al. (2020) | 24 | 7 | 77:23 | Passive surveillance Poland 2012-2018. Both positive and negative included. |
| **Total** | **928** | **631** | **60:40** | |

#### <u>References</u>

Amengual, B., Bourhy, H., López-Roig, M., & Serra-Cobo, J. (2007). Temporal dynamics of European bat Lyssavirus type 1 and survival of *Myotis myotis* bats in natural colonies. *PloS One*, 2(6), e566.

Barclay, R. M. (2012). Variable variation: annual and seasonal changes in offspring sex ratio in a bat. *PLoS One*, *7*(5), e36344.

Bat Conservation Trust (2022). Roost Count. Occurrence dataset https://doi.org/10.15468/vm4psx accessed via GBIF.org on 2023-01-12.

Blackwood, J. C., Streicker, D. G., Altizer, S., & Rohani, P. (2013). Resolving the roles of immunity, pathogenesis, and immigration for rabies persistence in vampire bats. *Proceedings of the National Academy of Sciences*, *110*(51), 20837-20842.

Broekman, M. J. E., Hoeks, S., Freriks, R., Langendoen, M. M., Runge, K. M., Savenco, E., ... & Tucker, M. A. (2023). HomeRange: A global database of mammalian home ranges. *Global Ecology and Biogeography*, *32*(2), 198-205.

Catto, C. M. C. (1993). Aspects of the ecology and behaviour of the serotine bat (*Eptesicus serotinus*), University of Aberdeen, UK.

Catto, C. M. C., Hutson, A. M., Raccey, P. A., & Stephenson, P. J. (1996). Foraging behaviour and habitat use of the serotine bat (*Eptesicus serotinus*) in southern England. *Journal of Zoology*, *238*(4), 623-633.

Chauvenet, A. L., Hutson, A. M., Smith, G. C., & Aegerter, J. N. (2014). Demographic variation in the UK serotine bat: filling gaps in knowledge for management. *Ecology and Evolution*, *4*(19), 3820-3829.

Colombi, D., Serra-Cobo, J., Métras, R., Apolloni, A., Poletto, C., López-Roig, M., ... & Colizza, V. (2019). Mechanisms for lyssavirus persistence in non-synanthropic bats in Europe: insights from a modelling study. *Scientific Reports*, *9*(1), 537.

Davis, A. D., Jarvis, J. A., Pouliott, C. E., Morgan, S. M., & Rudd, R. J. (2013). Susceptibility and pathogenesis of little brown bats (*Myotis lucifugus*) to heterologous and homologous rabies viruses. *Journal of virology*, *87*(16), 9008-9015.

Davis, A. D., Morgan, S. M., Dupuis, M., Poulliott, C. E., Jarvis, J. A., Franchini, R., … & Rudd, R. J. (2016). Overwintering of rabies virus in silver haired bats (*Lasionycteris noctivagans*). *PLoS One*, *11*(5), e0155542.

Fasel, N., Jeucken, J., Kravchenko, K., Fritze, M., Ruczynski, I., Komar, E., ... & Holtze, S. (2023). No intromission is involved in the mating of *Eptesicus serotinus*, a novel copulatory pattern in mammals.

Fleischer, T., Gampe, J., Scheuerlein, A., & Kerth, G. (2017). Rare catastrophic events drive population dynamics in a bat species with negligible senescence. *Scientific Reports*, *7*(1), 1-9.

Frick, W. F., Reynolds, D. S., & Kunz, T. H. (2010). Influence of climate and reproductive timing on demography of little brown myotis *Myotis lucifugus*. *Journal of animal ecology*, *79*(1), 128-136.

George, D. B., Webb, C. T., Farnsworth, M. L., O'Shea, T. J., Bowen, R. A., Smith, D. L., ... & Rupprecht, C. E. (2011). Host and viral ecology determine bat rabies seasonality and maintenance. *Proceedings of the National Academy of Sciences*, *108*(25), 10208-10213.

Glas, G. H. (1981). Activities of serotine bats (*Eptesicus serotinus*) in a nursing-roost. *Myotis*, *18*(19), 164.

Harbusch, C. (2003). *Aspects of the ecology of serotine bats (Eptesicus serotinus, Schreber 1774) in contrasting landscapes in southwest Germany and Luxembourg* (Doctoral dissertation, University of Aberdeen).

Harbusch, C., & Racey, P. A. (2006). The sessile serotine: the influence of roost temperature on philopatry and reproductive phenology of *Eptesicus serotinus* (Schreber, 1774)(Mammalia: Chiroptera). *Acta Chiropterologica*, *8*(1), 213-229.

Harris, S. L., Brookes, S. M., Jones, G., Hutson, A. M., & Fooks, A. R. (2006). Passive surveillance (1987 to 2004) of United Kingdom bats for European bat lyssaviruses. *Veterinary Record*, *159*(14), 439-446.

Kerth, G., & van Schaik, J. (2012). Causes and consequences of living in closed societies: lessons from a long-term socio-genetic study on Bechstein's bats. *Molecular ecology*, *21*(3), 633-646.

Kim, Y., Leopardi, S., Scaravelli, D., Zecchin, B., Priori, P., Festa, F., ... & Nouvellet, P. (2023). Transmission dynamics of lyssavirus in *Myotis myotis*: mechanistic modelling study based on longitudinal seroprevalence data. *Proceedings of the Royal Society B*, *290*(1997), 20230183.

Kurta, A., & Matson, J. O. (1980). Disproportionate sex ratio in the big brown bat (*Eptesicus fuscus*). *American Midland Naturalist*, 367-369.

Martinoli, A., Mazzamuto, M. V., & Spada, M. (2020). Serotine *Eptesicus serotinus* (Schreber, 1774). *Handbook of the Mammals of Europe*, 1-17.

Mathews, F., Kubasiewicz, L. M., Gurnell, J., Harrower, C. A., McDonald, R. A., & Shore, R. F. (2018). *A review of the population and conservation status of British mammals: technical summary*. Natural England.

Metheny, J. D., Kalcounis-Rueppell, M. C., Willis, C. K., Kolar, K. A., & Brigham, R. M. (2008). Genetic relationships between roost-mates in a fission–fusion society of tree-roosting big brown bats (*Eptesicus fuscus*). *Behavioural Ecology and Sociobiology*, *62*(7), 1043-1051.

Mills, R. S., Barrett, G. W., & Farrell, M. P. (1975). Population dynamics of the big brown bat (Eptesicus fuscus) in southwestern Ohio. *Journal of Mammalogy*, *56*(3), 591-604.

Moussy, C., Atterby, H., Griffiths, A. G. F., Allnutt, T. R., Mathews, F., Smith, G. C., ... & Hosken, D. J. (2015). Population genetic structure of serotine bats (*Eptesicus serotinus*) across Europe and implications for the potential spread of bat rabies (European bat lyssavirus EBLV-1). *Heredity*, *115*(1), 83-92.

Mühldorfer, K., Speck, S., Kurth, A., Lesnik, R., Freuling, C., Müller, T., ... & Wibbelt, G. (2011). Diseases and causes of death in European bats: dynamics in disease susceptibility and infection rates. *PLoS One*, *6*(12), e29773.

Orłowska, A., Smreczak, M., Freuling, C. M., Müller, T., Trębas, P., & Rola, J. (2020). Serological survey of lyssaviruses in polish bats in the frame of passive rabies surveillance using an enzyme-linked immunosorbent assay. *Viruses*, *12*(3), 271.

O'Shea, T. J., Ellison, L. E., Neubaum, D. J., Neubaum, M. A., Reynolds, C. A., & Bowen, R. A. (2010). Recruitment in a Colorado population of big brown bats: breeding probabilities, litter size, and first-year survival. *Journal of Mammalogy*, *91*(2), 418-428.

O'Shea, T. J., Ellison, L. E., & Stanley, T. R. (2011). Adult survival and population growth rate in Colorado big brown bats (*Eptesicus fuscus*). *Journal of Mammalogy*, *92*(2), 433-443.

Picard-Meyer, E., Barrat, J., Tissot, E., Verdot, A., Patron, C., Barrat, M. J., & Cliquet, F. (2006). Bat rabies surveillance in France, from 1989 through May 2005. *Developments in biologicals*, *125*, 283-288.

Racey, P. A., & Kleiman, D. G. (1970). Maintenance and breeding in captivity of some vespertilionid bats, with special reference to the noctule. *International Zoo Yearbook*, *10*(1), 65-70. [Cited in Harbusch, 2003- could not access]

Robardet, E., Borel, C., Moinet, M., Jouan, D., Wasniewski, M., Barrat, J., ... & Picard-Meyer, E. (2017). Longitudinal survey of two serotine bat (*Eptesicus serotinus*) maternity colonies exposed to EBLV-1 (European Bat Lyssavirus type 1): Assessment of survival and serological status variations using capture-recapture models. *PLoS Neglected Tropical Diseases*, *11*(11), e0006048.

Robinson, M. F., & Stebbings, R. E. (1997). Home range and habitat use by the serotine bat, *Eptesicus serotinus*, in England. *Journal of Zoology*, *243*(1), 117-136.

Shankar, V., Bowen, R. A., Davis, A. D., Rupprecht, C. E., & O'Shea, T. J. (2004). Rabies in a captive colony of big brown bats (*Eptesicus fuscus*). *Journal of wildlife diseases*, *40*(3), 403-413.

Stapelfeldt, B., Scheuerlein, A., Tress, C., Koch, R., Tress, J., & Kerth, G. (2022). Precipitation during two weeks in spring influences reproductive success of first-year females in the long-lived Natterer's bat. *Royal Society Open Science*, *9*(2), 211881.

Turbill, C., Bieber, C., & Ruf, T. (2011). Hibernation is associated with increased survival and the evolution of slow life histories among mammals. *Proceedings of the Royal Society B: Biological Sciences*, *278*(1723), 3355-3363.

Van der Poel, W. H., Van der Heide, R., Verstraten, E. R., Takumi, K., Lina, P. H., & Kramps, J. A. (2005). European bat lyssaviruses, the Netherlands. *Emerging Infectious Diseases*, *11*(12), 1854.

Vonhof, M. J., Strobeck, C., & Fenton, M. B. (2008). Genetic variation and population structure in big brown bats (*Eptesicus fuscus*): is female dispersal important? *Journal of Mammalogy*, *89*(6), 1411-1420.

Wilkinson, L. C., & Barclay, R. M. (1997). Differences in the foraging behaviour of male and female big brown bats (*Eptesicus fuscus*) during the reproductive period. *Ecoscience*, *4*(3), 279-285.

Wilkinson, G. S., & South, J. M. (2002). Life history, ecology, and longevity in bats. *Aging cell*, *1*(2), 124-131.

Zahn, A. (1999). Reproductive success, colony size, and roost temperature in attic-dwelling bat *Myotis myotis*. *Journal of Zoology*, *247*(2), 275-280.
