## Supplementary material for "Temporal variation in demography of temperate bats: consequences for population dynamics and disease": S2 Additional Figures

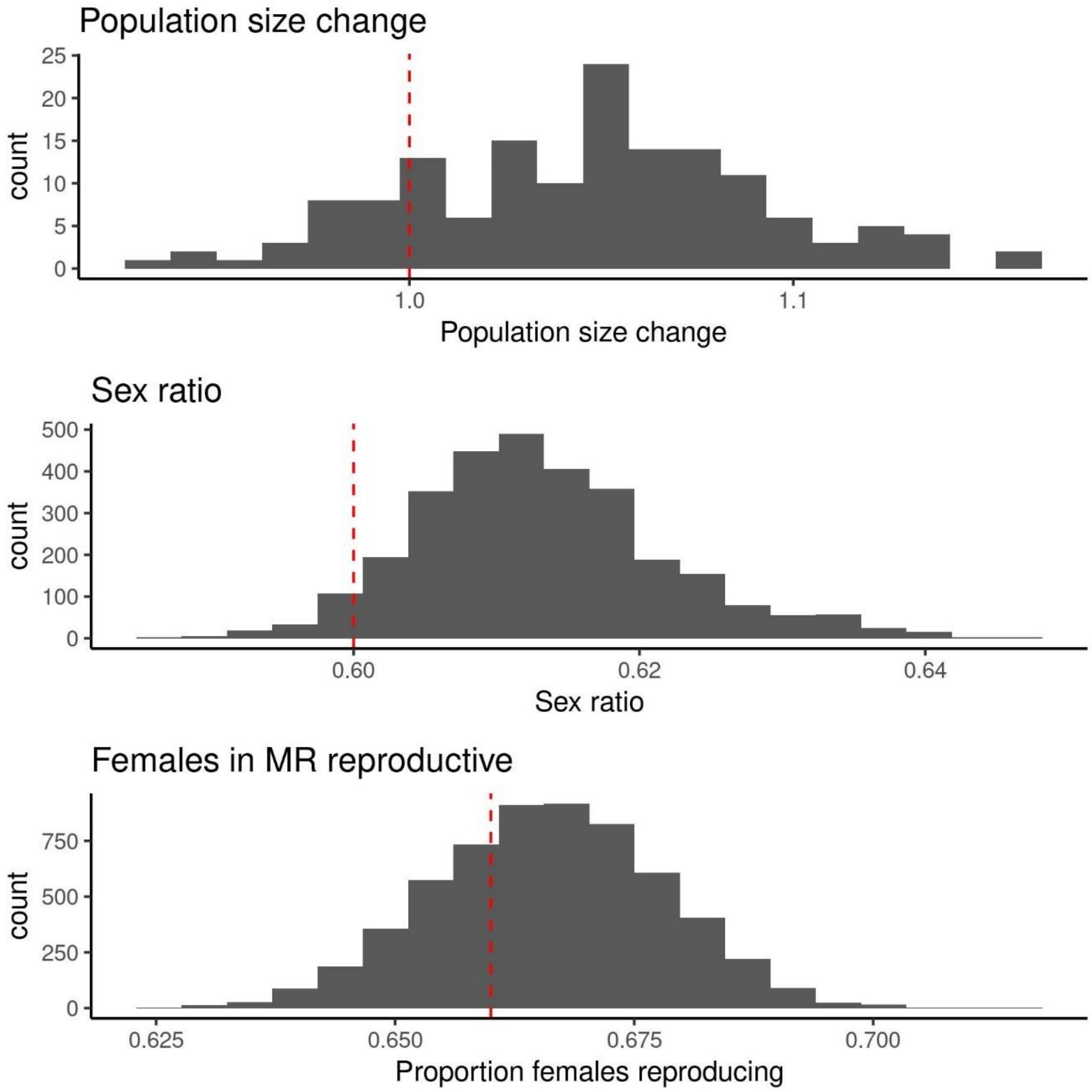


***Figure S1:*** *Base model fit relative to empirical observations. Red dotted lines indicate empirical estimates for population size change, sex ratio and the proportion of females reproducing within maternity roosts.*

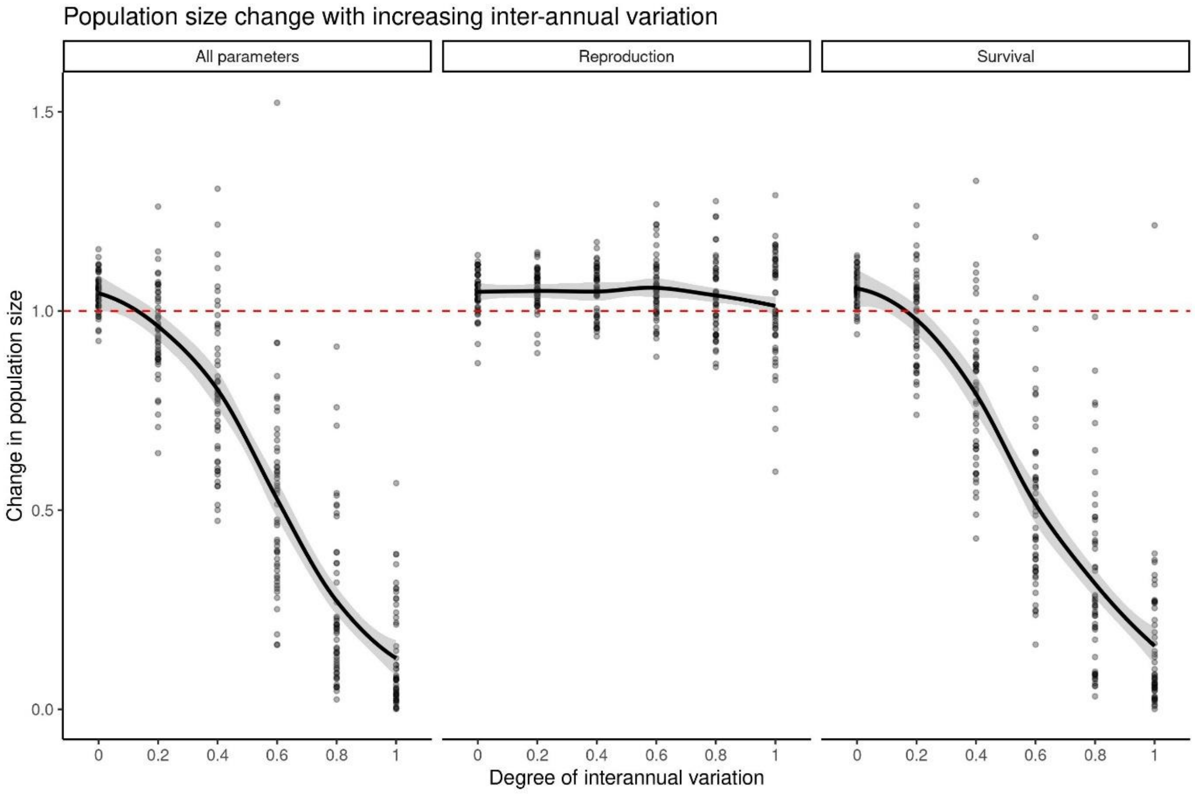


***Figure S2:*** *Influence of increasing inter-annual variation on population size when parameters are not correlated. This plot correlates to Figure 3, however in this case all parameters vary independently between years. Three scenarios were considered: all parameters vary between years, variation occurs only in reproductive parameters (primiparity, fecundity and pup survival) and variation occurs only in survival parameters (juvenile, non-breeder, and breeder survival). Trend lines were generated using the loess function from the geom_smooth function in R.*

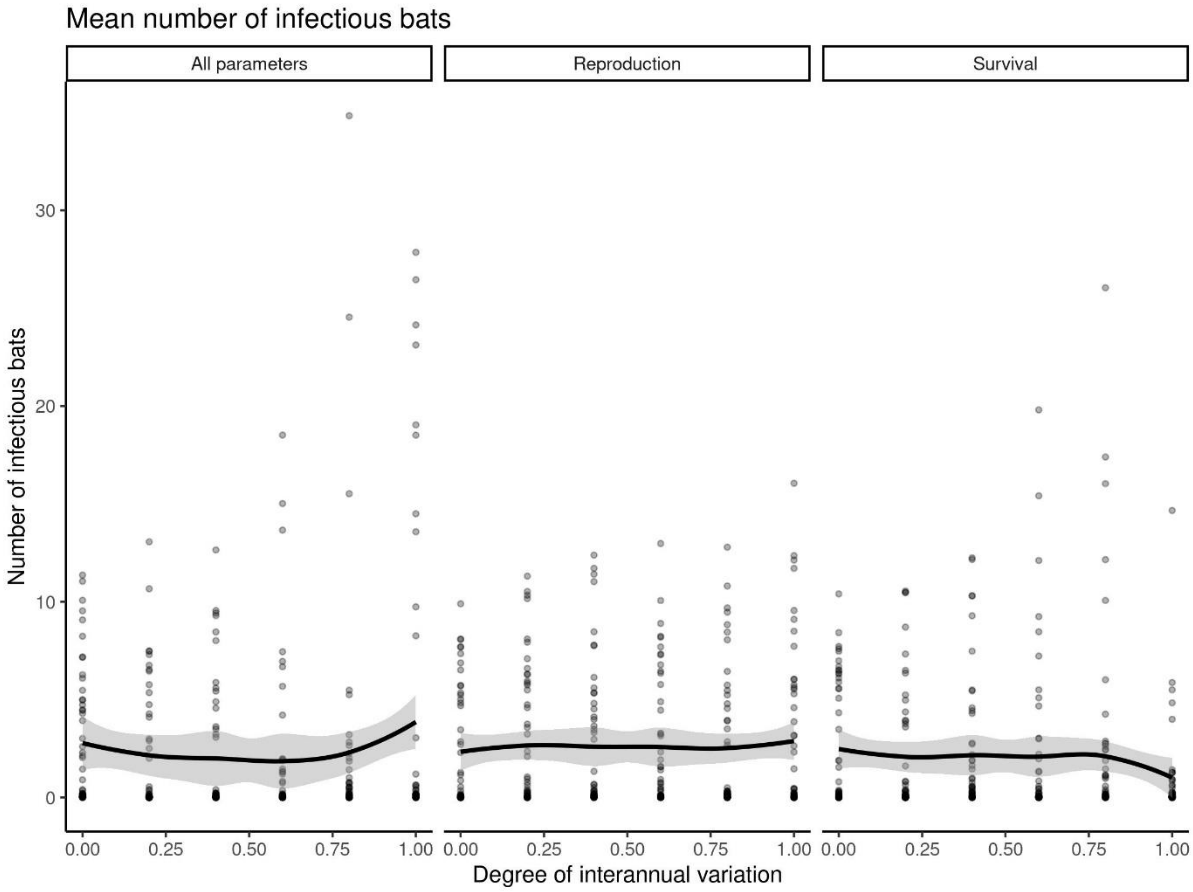


***Figure S3:*** *Influence of inter-annual variation on the total number of infected bats per year. Three scenarios were considered: all parameters co-vary between years, variation occurs only in reproductive parameters (primiparity, fecundity and pup survival) and variation occurs only in survival parameters (juvenile, non-breeder, and breeder survival). Trend lines were generated using the loess function from the geom_smooth function in R.*
